# Impacts of Disinfection Methods in a Granular Activated Carbon (GAC) Treatment System on Disinfected Drinking Water Toxicity and Antibiotic Resistance Induction Potential

**DOI:** 10.1101/2024.08.19.608195

**Authors:** Yinmei Feng, Stephanie S. Lau, William A. Mitch, Caroline Russell, Greg Pope, April Z. Gu

## Abstract

Graphic Abstract

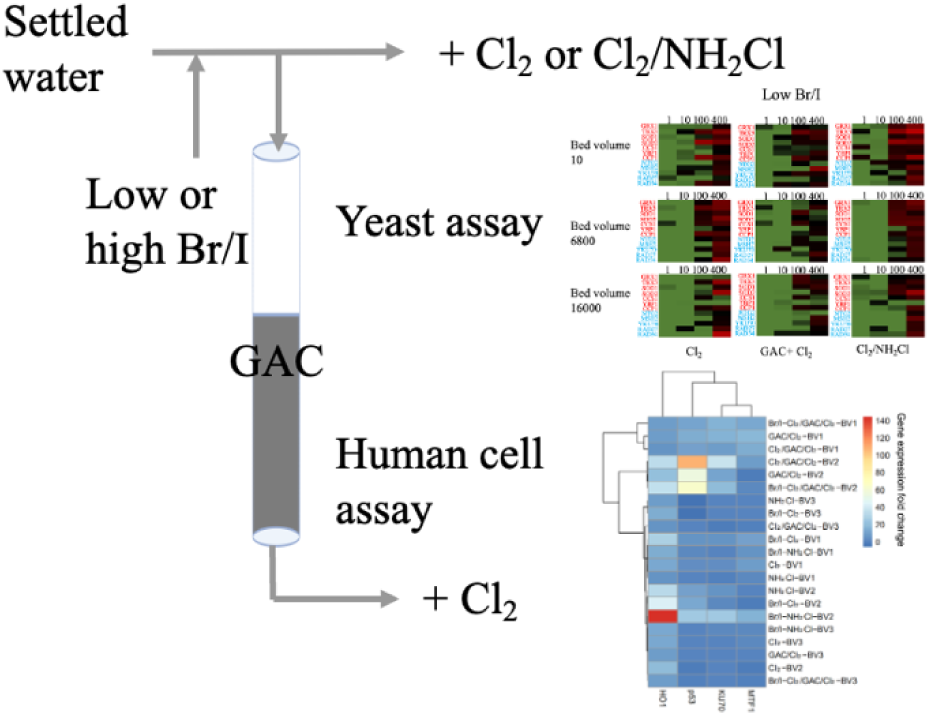

Granular activated carbon (GAC) treatment followed by chlorination (GAC/Cl_2_) and chlorination followed by chloramination (Cl_2_/NH_2_Cl) are two methods utilized by drinking water treatment facilities to mitigate the formation of disinfection byproducts (DBPs) in treated water. However, the effectiveness of these methods in reducing the overall toxicity of drinking water, driven by DBPs, remains largely unknown. In this study, we evaluate the total toxicity of water samples from a pilot-scale GAC system with post-chlorination (GAC/Cl_2_), and occasionally pre-chlorination upstream of GAC (Cl_2_/GAC/Cl_2_), compared to water treated by chlorination followed by chloramination (Cl_2_/NH_2_Cl). The research was conducted at various bromide and iodide levels and across three GAC bed volumes. To assess DNA stress and oxidative stress in water extracts, we employed the yeast toxicogenomic assay and human cell RT-qPCR assay, along with the DBP analysis from our previous study. Our results indicated that under environmental halogen conditions, GAC/Cl_2_ typically reduces both genotoxicity and oxidative stress in treated water more effectively than Cl_2_/NH_2_Cl and Cl_2_ treatment. However, Cl_2_/GAC/Cl_2_ does not consistently lower toxicity compared to GAC/Cl_2_. Notably, under high halogen conditions, Cl_2_/GAC/Cl_2_ fails to reduce genotoxicity and oxidative stress compared to samples without GAC treatment. Correlation analysis suggested that iodinated DBPs (I-DBPs) and nitrogenous DBPs (N-DBPs) were particularly associated with increased DNA stress and oxidative stress, indicating these classes of DBPs as significant contributors to the observed toxicity. While neither of these two categories of DBPs are regulated by the EPA, it appears that unregulated and unidentified DBPs significantly contribute to the genotoxicity and oxidative stress in drinking water. This research highlights the complex dynamics of water treatment processes and underscores the critical impact of unregulated DBPs on water toxicity.

## 1 Introduction

Chlorine disinfection a crucial process of drinking water treatment systems aimed at preventing waterborne diseases, however, epidemiological studies suggested an association between the exposure of chlorinated water and adverse human health outcomes, including risk of cancer and reproductive effects.(Cantor et al., 1998; Humans et al., 2004) Despite over 600 DBPs have been identified in drinking water, only 11 of them are regulated by Environmental Protection Agency (EPA), including four THMs (chloroform, bromodichloromethane, dibromochloromethane and bromoform) and five HAAs (monochloro-, monobromo-, dichloro-, dibromo-, and trichloroacetic acids).(Agency, 1979; Richardson, 2003) To ensure compliance with regulations, EPA has proposed utilizing alternative primary disinfectants such as ozone, UV radiation, or chlorine dioxide, adopting chloramines for residual maintenance in the distribution system, and implementing GAC to enhance the removal of total organic carbon (TOC) before primary disinfection as potential measures.(EPA, 2016)

Utilizing chloramines is an effective strategy for meeting residual requirements within the distribution system while complying with increasingly stringent regulatory limits for regulated DBPs. (Hua and Reckhow, 2008) A survey showed that 29% of the investigated water treatment facilities chose chloramine as secondary disinfection, since chloramination generally forms fewer THMs and HAAs than chlorination.(Seidel et al., 2005) On the other hand, other studies suggested that chloramination may induce the formation of unregulated nitrogenous disinfection byproducts (N-DBPs), although the total concentration of these compounds are relatively low.(Shah and Mitch, 2012) (Reckhow et al., 2001) Nonetheless, in vitro toxicological assays focusing on individual compounds have demonstrated that N-DBPs exhibit higher genotoxicity and cytotoxicity compared to the regulated THM and HAA species.(Richardson et al., 2007).

EPA considered GAC as one of the most promising technologies to control regulated DBPs, since it can efficiently remove the dissolved organic carbon (DOC), which serves as DBP precursors. (Chili et al., 2012; Summers et al., 1993) However, GAC is not effective at removing of halide ions and other inorganic DBP precursors, for example, bromide (Br^-^) leads to a higher Br to DOC ratio in the GAC effluent and might consequently result in a higher formation of Br-DBPs. (Chili et al., 2012; Erdem et al., 2020; Krasner et al., 2016; Summers et al., 1993; Symons et al., 1993) Moreover, it has been observed that GAC treatment demonstrates a preference for the removal of DOC rather than dissolved organic nitrogen (DON), and neither DON nor N-DBP precursors were effectively removed during the GAC treatment process.(Chili et al., 2012) Recent studies showed that Br-DBPs are generally more cyto- and geno-toxic than Cl-DBPs.(Jeong et al., 2015; Muellner et al., 2007; Plewa et al., 2004) Therefore, there is a concern that the benefits of GAC treatment in reducing the total concentration of measured DBPs may be offset by the formation of more toxic DBPs.

One approach to enhance GAC treatment performance is to use pre-chlorination to form DBPs within the treatment system prior to the GAC column, and then remove them using GAC or additional disinfection processes to eliminate precursors that lead to the formation of other DBPs.(Bond et al., 2012; Jiang et al., 2017) A recent study demonstrated that even with high concentration of Br-, GAC with pre-chlorination can remove the organic halogen efficiently and reduce the formation of regulated DBPs, however, there was another study showed a decrease formation of THMs, but no difference of HAAs after treated with GAC with pre-chlorination.(Fischer et al., 2019; Jiang et al., 2018) Furthermore, another study suggested that the tertiary-filtered wastewater with GAC treatment with pre-chlorination exhibited minimal or negligible effects on reducing DBP formation and calculated cyto- and geno-toxicity after the chlorination or chloramination of the GAC-treated effluent.(Verdugo et al., 2020) Due to the limit number of studies in this area, the removal rate of overall DBP formation might various with different GACs, source of natural organic matters and experimental conditions.(Erdem et al., 2020) Therefore, the assessment of various disinfection processes on DBP formation and their associated toxicity in drinking water, considering both low and high background levels of halogens, is of concern. In addition, recent studies revealed that DBPs in drinking water could also lead to antibiotic resistance by promoting genetic mutations and enhancing horizontal gene transfer.(Li and Gu, 2019; Zhang et al., 2017) These findings indicate that DBPs, particularly at sub-minimum inhibitory concentration levels, contribute to increased bacterial resistance through various mechanisms, including changes in cell membrane permeability, induction of SOS response, and altered conjugation-relevant gene expression.(Guo et al., 2015) Considering the complexity and synergistic effects of various DBPs in drinking water, drinking water may potentially induce antibiotic resistance.(Li and Gu, 2019) However, the impact of drinking water on the promotion of antibiotic resistance induction have hardly been evaluated.

Although the determination of calculated cytotoxicity and genotoxicity are valuable approaches to evaluate the quality of disinfected water, it is limited to approximately 100 measurable DBPs for which toxicity has been characterized using Chinese hamster ovary (CHO) cells.(Wagner and Plewa, 2017) Recent studies suggested that these known DBPs may not be the main toxic drivers in the disinfected waters. (Lau et al., 2023a; Wu et al., 2020) Great knowledge gaps remain regarding the toxicological implications of the combined effects of both known and unknown DBPs on the overall quality of disinfected water. However, traditional animal-based toxicological studies are time-consuming and costly, while single endpoint in vitro assays have limitations in comprehensive evaluation, indicating the importance of developing a toxicogenomics-based rapid screening approach for drinking water toxicity analysis to enhance mechanistic understanding and identify major toxicity drivers.(Neale et al., 2012; Shukla et al., 2010) Furthermore, as we discussed previously, the potential of drinking water containing a mixture of DBPs on antibiotic resistance induction have never been investigated. The objective of this study was to 1) investigate the impact of GAC treatment with and without pre-chlorination on DBP formation and the resulting genotoxicity of drinking water samples, 2) Use genotoxicity toxicogenomics bioassays to provide a profiling and quantitative analysis of toxicity across the drinking water samples with or without pre-chlorination. 3) Apply a newly proposed toxicogenomics assay to evaluate the antibiotic resistance induction potential of the disinfected drinking water samples.

In this study, effluents from a pilot-scale GAC system with post-chlorination, and occasionally pre-chlorination upstream of GAC, as well as the GAC influents treated by chlorine followed by chloramine and chlorine, were analyzed. We performed the genotoxicity and antibiotic resistance potential assessment. Our results demonstrated that GAC treatments with post-chlorination can enhance the reduction of genotoxic compounds and decrease the induction potential for antibiotic resistance, while GAC with pre-chlorination cannot consistently reduce the genotoxicity of treated water compared to GAC without pre-chlorination. These results highlighting the effectiveness of GAC in improving water quality and safety.

## 2 Material and Methods

### 2.1 GAC pilot system

The details of the GAC pilot system was described in our previous study, and in SI.(Lau et al., 2023b) Briefly, three parallel 6.4-cm diameter granular activated carbon (GAC) columns were operated to evaluate different conditions (Figure 1) including Column 1: GAC treatment without bromide (Br^-^) and iodide (I^-^) spiking or pre-chlorination, Column 2: GAC treatment without Br- and I-spiking but with pre-chlorination, and Column 3: GAC treatment with Br^-^ and I^-^ spiking and pre-chlorination.

**Figure 1:**
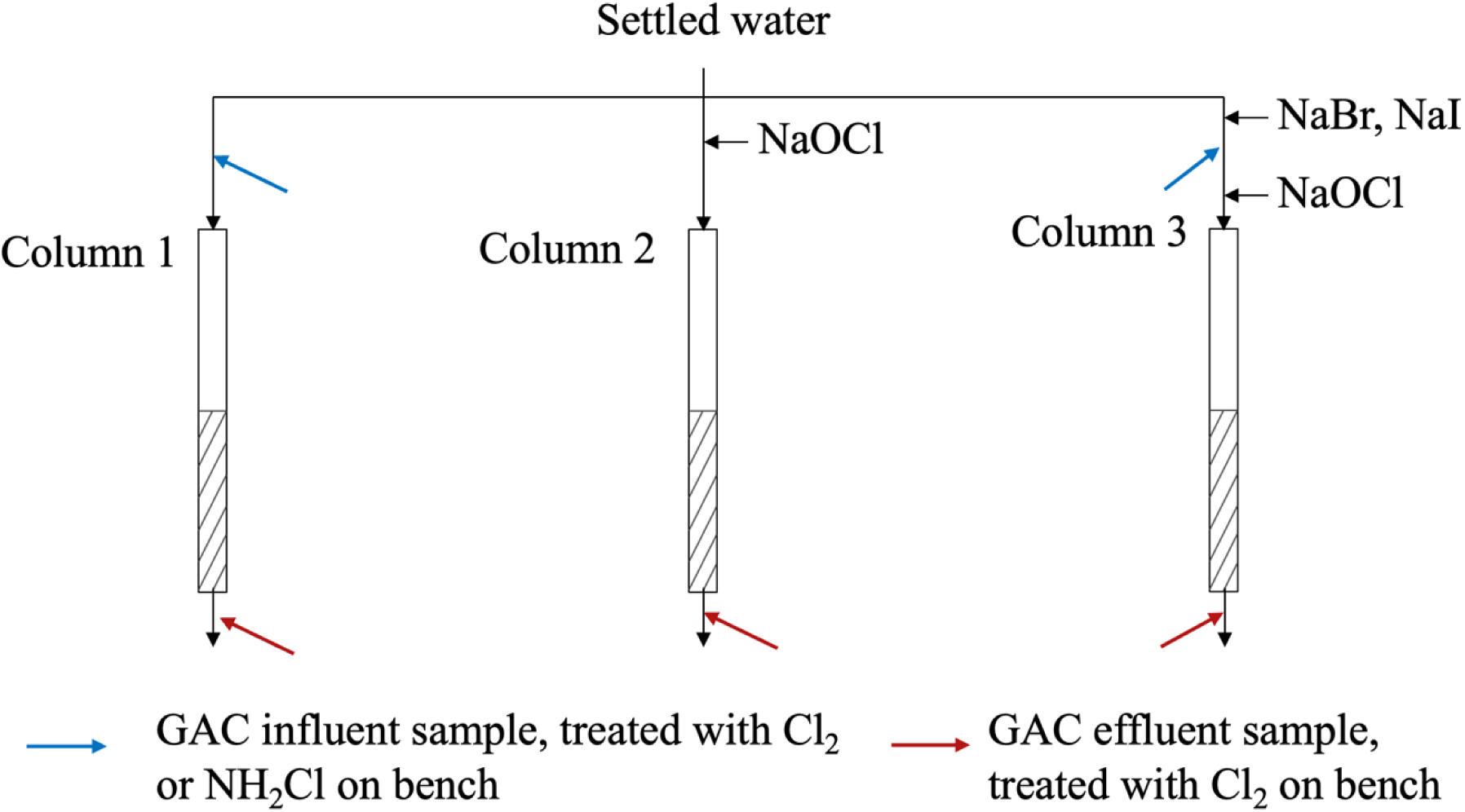
Schematic of the GAC pilot system.

Water samples were collected during three sampling events in September 2021, December 2021, and February 2022, corresponding to 10, 6800, and 16000 bed volumes (BV) of water treated through the GAC system; analysis of TOC measurements upstream and downstream of the GAC columns revealed breakthrough percentages of approximately 20-30%, 40-45%, and 60-65% for the respective sampling events.

### 2.2 Sample preparation and extraction

Water samples from the GAC system were treated with chlorine or chlorine/chloramines and then quenched with sodium thiosulfate after a 3-day contact period. These quenched samples were acidified and processed using custom-made solid-phase extraction cartridges. The cartridges, containing 2.5 g of Sepra ZTL sorbent, were pre-conditioned and equilibrated before loading the samples. The water samples were then extracted through the cartridges, followed by drying, elution with methanol, and concentration of the eluates. The final extracts were prepared using a DMSO-methanol solvent exchange method. Detailed method is described in the Text S2.(Lau et al., 2023b)

### 2.3 Quantitative toxicogenomics assay for genotoxicity assessment

A quantitative high-throughput toxicogenomics-based GFP-fused yeast reporter library assay was employed to fast and mechanistically evaluate the toxicity of collected water samples with high sensitivity as described in our previous publications.(Lan et al., 2018) In brief, a library of yeast reporters with in-frame green florescent protein (GFP) fusion of *Saccharomyces cerevisiae* (Invitrogen, no. 95702, ATCC 201388) were constructed by oligonucleotide-directed homologous recombination to tag the open reading frames (ORF) with Aequorea victoria GFP (S65T) in its chromosomal location at the 3′ end. In this study, twelve biomarkers (yeast reporter strains) were selected based on their high correlations with and predictability of phenotypic endpoints of genotoxicity and oxidative stress, selected biomarkers can be found in SI. The selected yeast strains were seeded in SD medium with Dropout (DO) supplement-His (Clontech, CA, US) on a sterile black clear-bottom 384-well plate (Corning, USA) and incubated for 4-6 h at 30 °C to reach the early exponential phase. 10 μL of water extracts diluted with PBS, blank control (SD medium + 0.25 YPD medium, with and without chemical), and internal housekeeping control (SD medium + 0.25 YPD medium + PGK1 strain, with and without chemical) were added to the designated wells to achieve the intended concentrations. The absorbance (600 nm) and fluorescence signals (excitation at 485 nm and emission at 535 nm) of test wells were measured simultaneously using a Microplate Reader (Synergy H1 Multi-Mode, Biotech, Winooski, VT) every 5 min for 2 h. The expression levels of the biomarkers and cell growth were reflected by the GFP and OD signals, respectively. All experiments were performed in triplicate in the dark.

Data processing of protein expression profiling and derivation of molecular toxicity endpoints were previously described(Lan et al., 2018). The molecular toxicity endpoint, Protein Effect Level Index (PELI) quantifies the accumulative altered protein expression change over the 2-hour exposure period for a given protein (*PELI_ORFi_*) or a given stress category (*PELI_geno_* and *PELI_oxi_* for DNA and oxidative stress, respectively). The pathway activation response is calculated by the following equation:

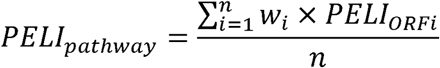

where n represent the number of ORFs in the pathway, and w_i_ is the weight factor of ORFi. In this study, a value of 1 were assigned for all weight factors.

A cutoff threshold PELI value of 1.5 was determined to represent a statistically significant increase in protein expression levels compared to the untreated control based on previous control experiments, the standard deviation range of our testing systems, and the commonly used threshold of fold change in protein expression in literature. For each water sample, dose-response relationships of both *PELI_geno_*and *PELI_oxi_* were fitted using logistic regression model in Prism 9. Molecular toxicity endpoint PELI1.5 was defined as the sample concentrations that corresponded to a PELI value of 1.5. The DNA and oxidative stress of each water sample were expressed as toxic equivalents of reference compounds as described previously:

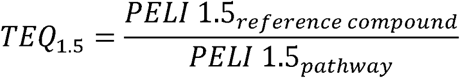

where, 4-nitroquinoline 1-oxide (4-NQO) was used as reference compound for genotoxicity(Martijn et al., 2016) and hydrogen peroxide (H2O2) was used as reference compound for oxidative stress(Jiang et al., 2021).

### 2.4 Human cell RT-qPCR assay

Reverse transcription quantitative PCR (RT-qPCR) technology was employed to quantify the relative transcription level changes of 4 key genes indicative of four cellular stress and toxicity response pathways, including DNA damage (*KU70*), oxidative stress (*HO1*), chemical stress (*MTF-1*) and apoptosis (*Casp3*) in exposure to collected water samples with a human liver epithelial cell line (HepG2). Approximately 2×10^5^ per well of HepG2 cells were seeded in Eagle’s Minimum Essential Medium (ATCC) with 10% fetal bovine serum (GE Healthcare Life Sciences, US) on a sterile 6-well plate (Corning) and cultured at 37°C for 24 h. 1 ml of water samples diluted with EMEM medium were added in each well. After 4 h of exposure, RNA from exposed cell samples were extracted using one-step RNA reagent (Bio Basic Inc., Canada), and reverse transcribed to cDNA using Verso cDNA Synthesis Kit (Thermo Scientific, U.S.A). Q-PCR was performed in triplicate using SYBR Green Supermix (Bio-Rad, U.S.A.) on iQ7 Multicolor Real-Time PCR detection system (Bio-Rad, U.S.A.). PCR primers targeting the selected stress genes were obtained based on literature review and the NCBI database as previously described.(Lin et al., 2020) Housekeeping gene 18s rRNA was used as internal control to normalize the quantities of the target genes. Alterations of gene expression levels in exposure to water samples compared to untreated control were expressed as fold changes, also referred to as induction actor *I*, determined by the comparative C_T_ (2^−ΔΔCT^) method as reported in literature.(Schmittgen and Livak, 2008)

### 2.5 Transcriptional Analysis of Selected Biomarkers in *E. coli* Cells for Antibiotic resistance potential assessment

A library of 18 transcriptional fusions of GFP that includes promoters controlling the expression of genes involved in outer membrane permeability, efflux pump, drug inactivation, targets alternations, SOS response and detoxifications in E. coli K12, MG1655 was employed in this study. The rationale for the biomarker selection, their roles involved in antibiotic resistance are described in SI.(Gou et al., 2014) In brief, GFP-fused E. coli strains selected were grown with M9 medium in clear bottom black 384-well plates for 4–6 h at 37 °C to reach early exponential growth stage (OD600 value of 0.15-0.25). Freshly prepared water samples or PBS control were added at CF=100 per well. A microplate reader was used to read plates for absorbance (OD600 for cell growth) and GFP signal (filters with 485 nm excitation and 535 nm emission for gene expression) every 5 min for 2 h. (Lan et al., 2014)

Data processing of protein expression profiling and derivation of molecular toxicity endpoints can be found on SI. The molecular antibiotic resistance induction potential index (ARIPI) was used to quantify the gene expression change over the 2-hour exposure period.

### 2.6 Statistical analysis

Maximum Cumulative Ratio (MCR) calculation was employed to assess the health risks and prioritize contaminants in the disinfected water samples. In the calculation of hazard quotients (HQs), for TTHMs and HAA5, which are regulated in drinking water by EPA, their permitted doses were selected as regulatory maximum contaminant level (MCL). For other non-regulated chemicals, the LC50 data from a CHO cell chronic cytotoxicity assay reported in literature were used as MCLs.(Wagner and Plewa, 2017) The detailed calculation can be found in SI.

Hierarchical clustering analysis was performed with complete linkage. Spearmen correlation was then carried out in R to identify the potential relationships between the molecular toxicity quantifiers and DBP concentrations in drinking water samples at concentration factor of 100.

## 3 Results and discussion

### 3.1 Multi-toxicity assay with both yeast and human cells revealed the impact of disinfection process on drinking water toxicity

#### 3.1.1 The impact of disinfection process on drinking water toxicity profiles

To assess the impact of different water disinfection processes on drinking water toxicity, organic extracts from 21 water samples collected from a pilot-scale GAC system at varying BVs were analyzed using a toxicogenomic-based yeast assay. This study revealed distinct molecular toxicity fingerprints represented by the PELI, which targeted 12 key biomarkers related to oxidative and DNA stress. Overall, the cellular toxicity patterns observed in these samples are concentration-dependent and treatment process dependent, with elevated protein expression levels in both DNA and oxidative stress revealed at higher concentration factor (CF). Further comparative analysis using toxic equivalents (geno-TEQ1.5 and oxi-TEQ1.5) allowed for a more detailed assessment of the differential impact of each disinfection process on these cellular stress pathways, with higher TEQ value represent higher toxicity of the sample.

##### 3.1.1.1 Contribution of halogens to the toxicity of drinking water disinfected with chlorine (Cl_2_) versus those disinfected with the combination of chlorine and chloramine (Cl_2_/NH_2_Cl)

Halogen is an important class of inorganic DBP precursor in drinking water, and the concentration of halogens in the source water can influence the toxicity of treated drinking water. Therefore, we assessed the toxicity of drinking water treated with either chlorine or chloramine at various levels of bromine and iodine concentration.

In samples collected from the influents of the GAC system at different BVs, Cl_2_ generally reduced overall toxicity in most samples, as compared to treatments with Cl_2_/NH_2_Cl (Figure 2a). Under environmental background level of halogens combined with Cl_2_ treatment, no biomarkers were upregulated in drinking water extracts at relatively low concentration factors (CF=1, 10).

**Figure 2:**
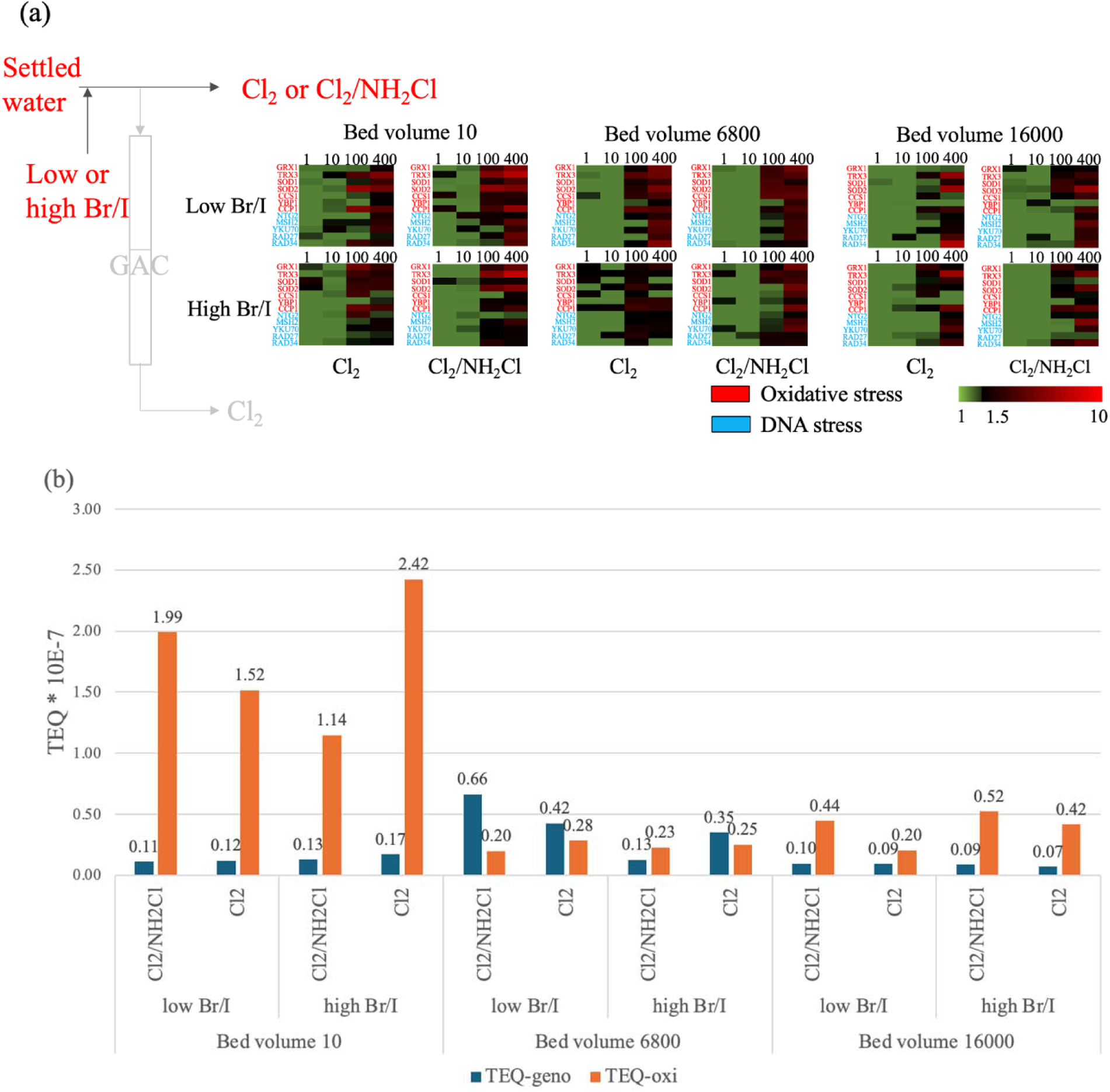
(a) Altered protein expression profiles of 12 biomarkers indicative of oxidative stress and DNA stress responses upon exposure to GAC influents treated with free chlorine only (Cl_2_) and chlorine/chloramines (Cl_2_/NH_2_Cl) from GAC system at different BVs. PELI values are scaled by the green-black-red color spectrum at the bottom right. Green to black color bars indicate no significant protein expression level changes, and black to red color bars indicate elevated up-regulation. X-axis top: concentration of the water samples in units of concentration factors. X-axis bottom: treatment process. Y-axis right: list of protein biomarkers color-coded based on their associated stress categories (red: oxidative stress; blue: DNA stress) (b) Summary of integrated molecular toxicity endpoints (TEQgeno and TEQoxi) for DNA and oxidative stress of GAC influents treated with free chlorine only (Cl_2_) and chlorine/chloramines (Cl_2_/NH_2_Cl) from GAC system at different BVs.

Conversely, low CF of samples treated with Cl_2_/NH_2_Cl exhibited upregulation of Cytochrome C (CCP1), superoxide dismutase (CCS1), and base excision repair (NTG2) related biomarkers. At higher concentrations (CF=100 and 400), samples with environmental background of halogens treated with Cl_2_/NH_2_Cl appeared more toxic than those treated with Cl_2._ (Figure S1) This increased toxicity from Cl_2_/NH_2_Cl may be due to the formation of N-DBPs, which tend to be more toxic than carbonaceous DBPs (C-DBPs). Similarly, in samples with high halogen level, those treated with Cl_2_/NH_2_Cl exhibited increased toxicity at high CF, while no significant differences were observed at low CF.

We then compared the quantitative endpoint of samples treated with Cl_2_ versus Cl_2_/NH_2_Cl at different halogen level. In consistent with the discussion earlier, GAC influents with low halogen level, samples treated with Cl_2_/NH_2_Cl generally lead to higher DNA stress than Cl_2_ ones, except the BV 10 batch, this trend was similar with the additive cytotoxicity comes from unregulated DBPs. As for oxidative stress, samples treated with Cl_2_/NH_2_Cl also induce higher toxicity than Cl_2_ ones, except the BV 6800 batch. Despite these exceptions, the difference in DNA and oxidative stress between these two treatments were minimal. For GAC influents with high halogen level, Cl_2_/NH_2_Cl treatment may lead to the reduction on both oxidative and DNA stress, compared to Cl_2_, for BV 10 and 6800, but not for BV 16000. (Figure 2b) While this result is similar with the previous total cytotoxicity result, the oxidative stress was also generally higher in the high halogen waters than in the low halogen ones. The potential reason behind this increase may related to the chlorination of Br contained water leads to the formation of HOBr, which exhibits a higher reactivity towards organic compounds compared to HOCl. (Lee and Gunten, 2009; Li et al., 2020)

##### 3.1.1.2 Impact of GAC with post-chlorination (GAC/Cl_2_) versus combination of chlorination and chloramination (Cl_2_/NH_2_Cl) on the toxicity of drinking water

To comply with the EPA’s regulations on DBPs, employing GAC treatment prior to disinfection process, or using Cl_2_/NH_2_Cl as disinfectants, are two common strategies to reduce the formation of regulated DBPs. We assessed the toxicity of GAC effluent treated with chlorine versus GAC influent treated with Cl_2_/NH_2_Cl. As shown in Figure 3a, GAC/Cl_2_ generally reduce both oxidative and DNA stress of water samples compared to GAC influent treated with Cl_2_ or Cl_2_/NH_2_Cl. At high concentration (CF=100, 400), the PELI values of DNA stress of the GAC-treated samples were lower than those without GAC treatment. For example, NTG2, a base pair repair biomarker, only show an upregulation (PELI>1.5) in the samples without GAC treatment. At low CF, no altered protein expression was observed for any of the selected biomarkers. (Figure S2).

**Figure 3:**
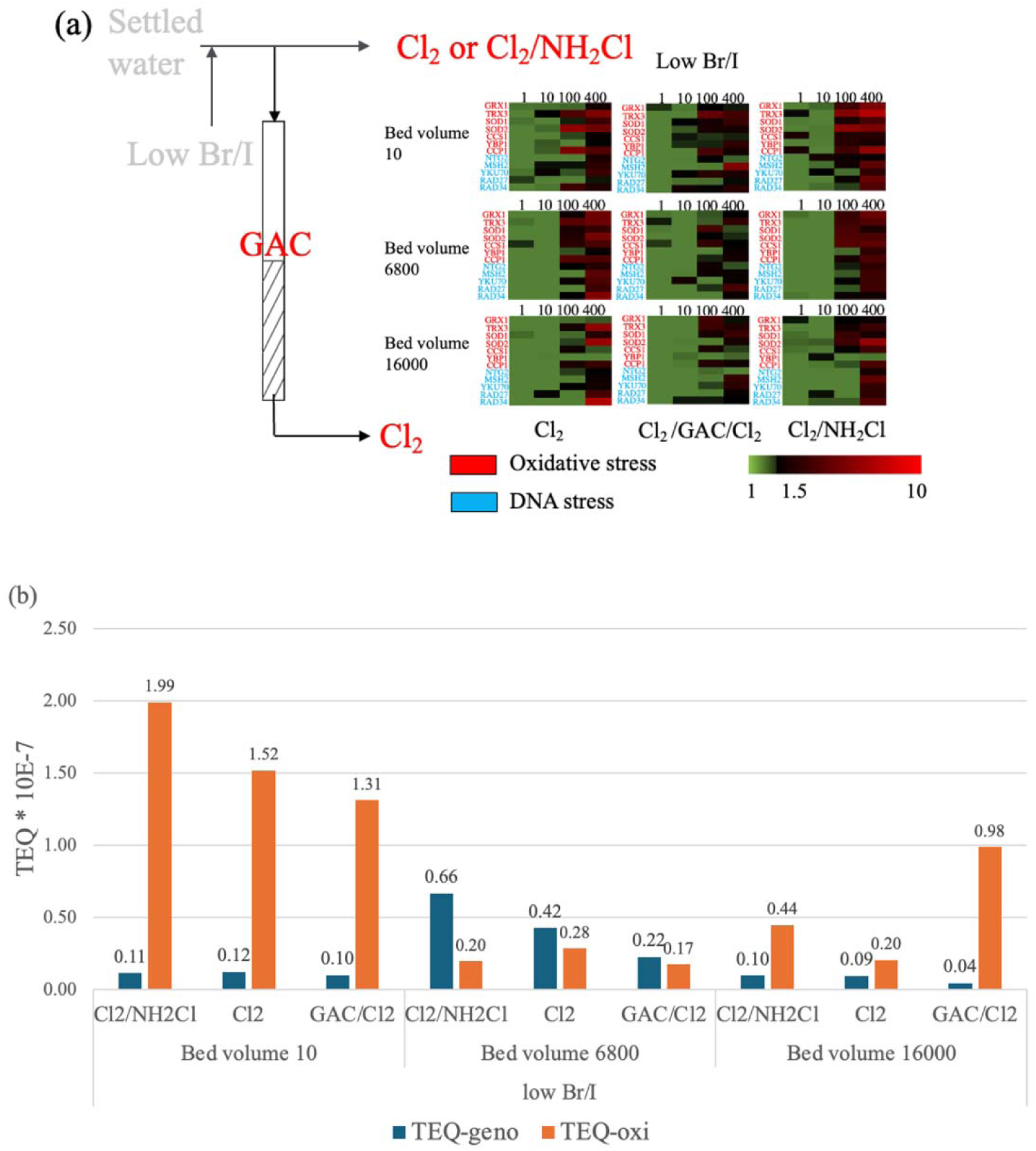
(a) Altered protein expression profiles of 12 biomarkers indicative of oxidative stress and DNA stress responses upon exposure to water samples with low halogen that were treated by chlorination (Cl_2_), GAC with post-chlorination (GAC/Cl_2_), or chlorination/chloramination (Cl_2_/NH_2_Cl) from GAC system at different BVs. PELI values are scaled by the green-black-red color spectrum at the bottom right. Green to black color bars indicate no significant protein expression level changes, and black to red color bars indicate elevated up-regulation. X-axis top: concentration of the water samples in units of concentration factors. X-axis bottom: treatment process. Y-axis right: list of protein biomarkers color-coded based on their associated stress categories (red: oxidative stress; blue: DNA stress) (b) Summary of integrated molecular toxicity endpoints (TEQgeno and TEQoxi) for DNA and oxidative stress of water samples with low halogen level that were treated by chlorination (Cl_2_), GAC with post-chlorination (GAC/Cl_2_), or chlorination/chloramination (Cl_2_/NH_2_Cl) from GAC system at different BV.

We then compared the oxidative stress and DNA stress endpoint of drinking water samples with GAC/Cl_2_ and with Cl_2_/NH_2_Cl under low halogen level. Our results showed that the genotoxicity was lower in GAC/Cl_2_ treatment samples, under all three BVs, (Figure 3b) which is similar with our previous results that chloramination was less effective than GAC for reducing the CHO cell cytotoxicity with known DBPs.(Lau et al., 2023b) GAC treatment seemed to reduce both genotoxicity and oxidative stress in most of the samples with low halide concentration (except the oxidative stress at BV 16000, this may be explained by the less effective of GAC with the increasing of its lifetime).(Peterson et al., 2022) Our previous study also demonstrated that the removal efficient of GAC on the DBP formation reduced with the increase of GAC lifetime.(Lau et al., 2023b)

##### 3.1.1.3 Impact of pre-chlorination on the GAC performance in the reduction of drinking water toxicity

To improve the efficiency of GAC in removing DBPs, some studies have suggested that incorporating an additional chlorination step before the GAC treatment (Cl_2_/GAC/Cl_2_), enabling direct removal of DBPs rather than their precursors, can reduce the concentration of regulated DBPs. However, other research argued that in practical drinking water treatment processes, this could lead to increased overall DBP formation and result in more toxic treated water. Our results indicate that under low halogen level, GAC appears to reduce toxicity compared to samples without GAC treatment. (Figure 3). Yet, no consistent differences were observed across all concentration factors between samples treated with Cl_2_/GAC/Cl_2_ and GAC/Cl_2_. At low concentration (CF=1,10), samples treated with Cl_2_/GAC/Cl_2_ exhibited higher toxicity, particularly impacting RAD27 (base excision repair related) and MSH2 (mismatch repair related) biomarkers. At high concentration (CF=100,400), no significant differences were identified between samples treated with Cl_2_/GAC/Cl_2_ and GAC/Cl_2_. (Figure 4a, Figure S2)

**Figure 4:**
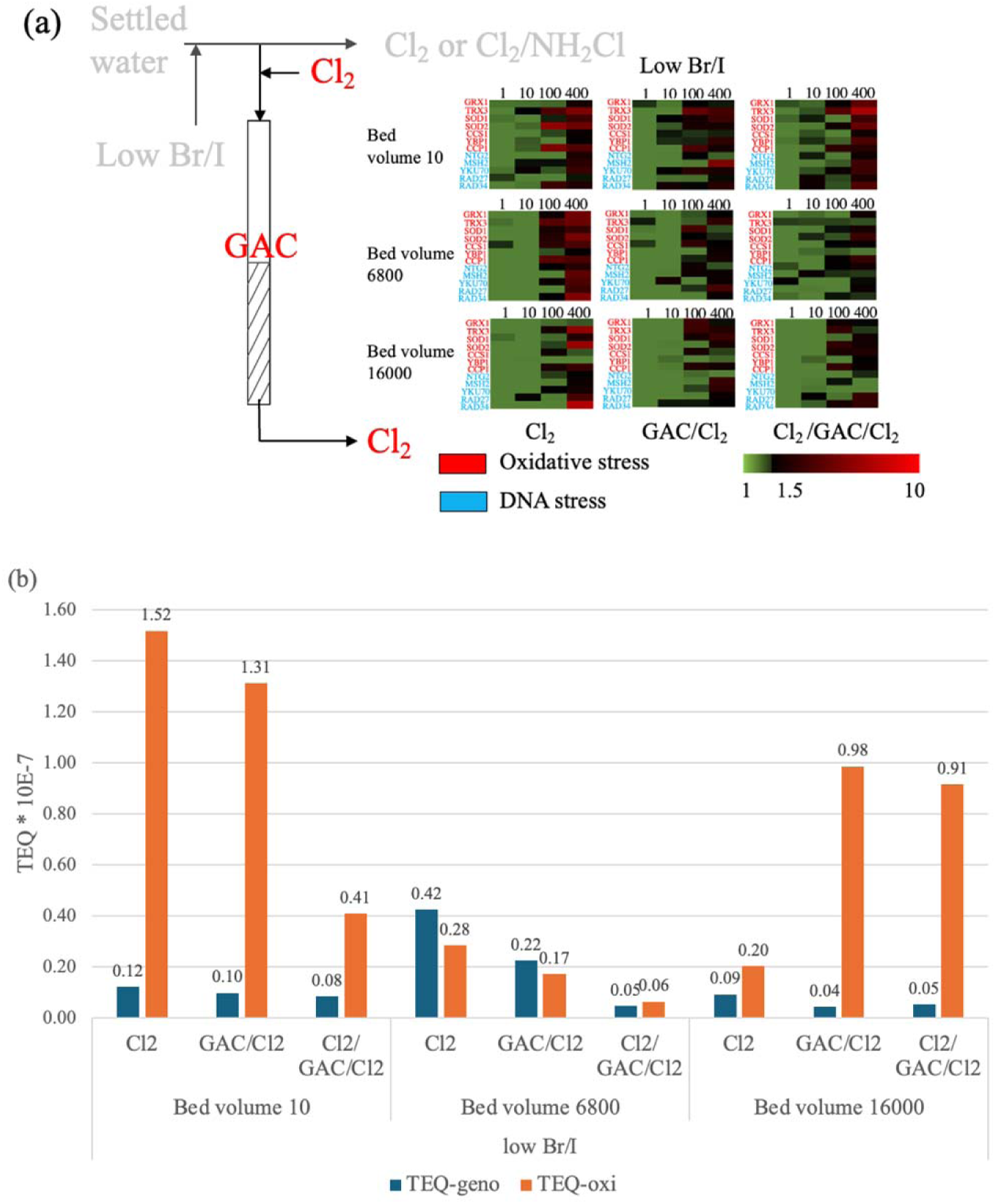
(a) Altered protein expression profiles of 12 biomarkers indicative of oxidative stress and DNA stress responses upon exposure to water samples with low halogen that were treated by chlorination (Cl_2_), GAC with post-chlorination (GAC/Cl_2_), or GAC with pre- and post-chlorination (Cl_2_/GAC/Cl_2_) from GAC system at different BVs. PELI values are scaled by the green-black-red color spectrum at the bottom right. Green to black color bars indicate no significant protein expression level changes, and black to red color bars indicate elevated up-regulation. X-axis top: concentration of the water samples in units of concentration factors. X-axis bottom: treatment process. Y-axis right: list of protein biomarkers color-coded based on their associated stress categories (red: oxidative stress; blue: DNA stress) (b) Summary of integrated molecular toxicity endpoints (TEQgeno and TEQoxi) for DNA and oxidative stress of water samples with low halogen that were treated by chlorination (Cl_2_), GAC with post-chlorination (GAC/Cl_2_), or GAC with pre- and post-chlorination (Cl_2_/GAC/Cl_2_) from GAC system at different BVs.

We evaluated the performance of GAC/Cl_2_ and Cl_2_/GAC/Cl_2_ across different BVs in the treatment of source water with low halide level, by using the TEQ1.5. Figure 4b illustrates that for water with low halogen level, Cl_2_/GAC/Cl_2_ generally reduced both genotoxicity and oxidative stress more effectively than GAC/Cl_2_, except at the BV of 16000. Although the differences in DNA stress and oxidative stress under the treatment between GAC/Cl_2_ and Cl_2_/GAC/Cl_2_ were minor at BVs 10 and 16000, a large reduction occurred at BV 6800. Notably, this reduction did not correspond with the total cytotoxicity results, which suggested similar levels of cytotoxicity removal of GAC/Cl_2_ or Cl_2_/GAC/Cl_2_ across three BVs. However, at BV 6800, a 32.5% reduction in the calculated cytotoxicity of known unregulated DBPs was observed with Cl_2_/GAC/Cl_2_ compared to GAC/Cl_2_. This aligns with the substantial reductions in DNA and oxidative stress, suggesting that the toxicity of these samples may be predominantly influenced by unregulated DBPs.

##### 3.1.1.4 Effect of GAC with pre-chlorination (Cl_2_/GAC/Cl_2_) versus chloramination (Cl_2_/NH_2_Cl) under high level of halide

We further assessed the performance of Cl_2_/GAC/Cl_2_ in reducing the overall toxicity of water samples under high halogen level. However, as indicated in Figure 5a under high halogen concentration, Cl_2_/GAC/Cl_2_ failed to diminish the toxicity of the water samples. Figure S2 further demonstrates that there is no reduction in the disruption of the PELI for selected biomarkers between samples treated with Cl_2_/GAC/Cl_2_ and without GAC. These findings suggest that while GAC treatment generally reduces the overall toxicity of drinking water samples, introducing pre-chlorination to the treatment process at high halogen concentrations may not be beneficial and can sometimes even impair GAC performance.

**Figure 5:**
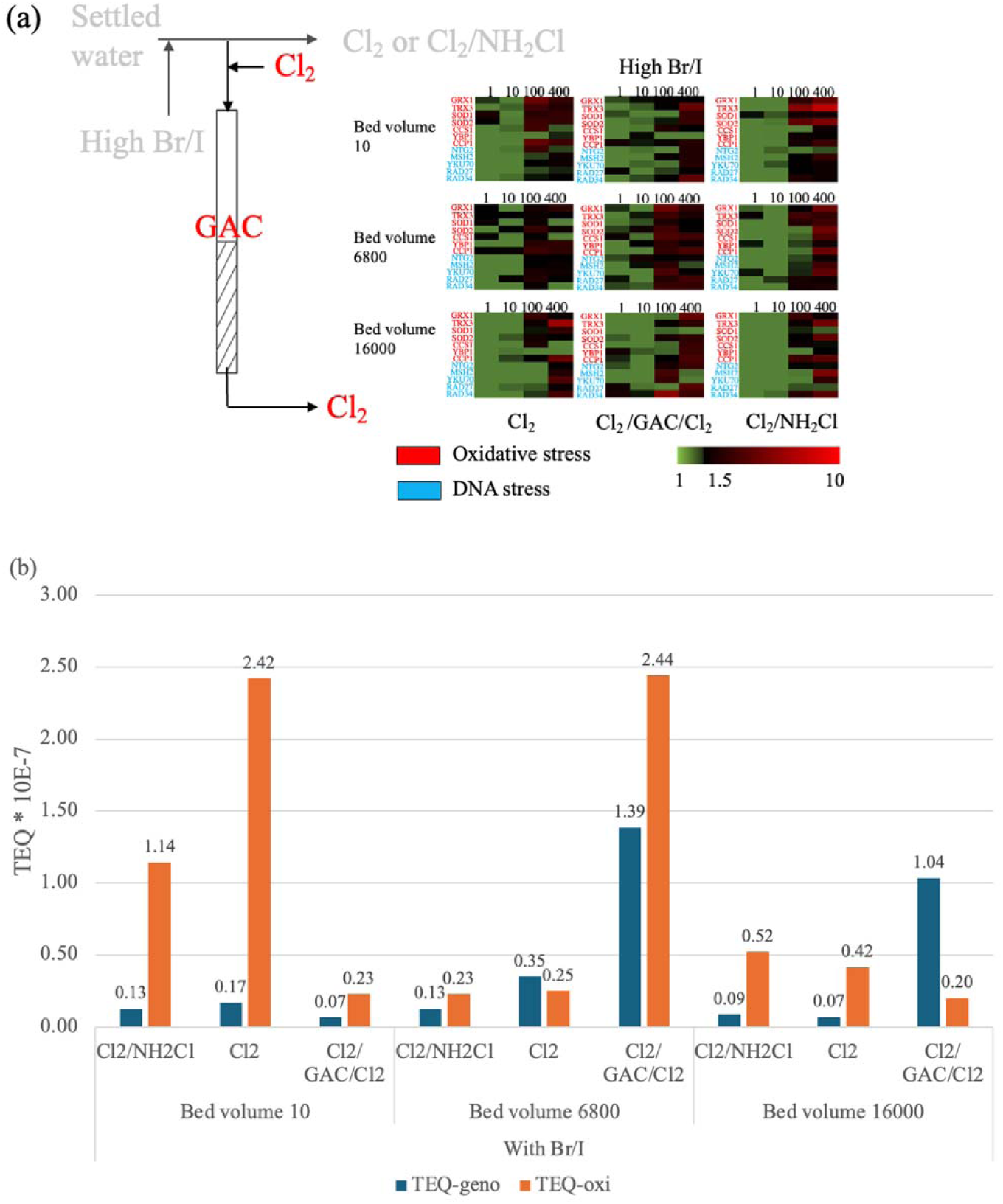
(a) Altered protein expression profiles of 12 biomarkers indicative of oxidative stress and DNA stress responses upon exposure to water samples with high halogen that were treated by chlorination (Cl_2_), chlorine/chloramines (Cl_2_/NH_2_Cl), or GAC with pre- and post-chlorination (Cl_2_/GAC/Cl_2_) from GAC system at different BVs. PELI values are scaled by the green-black-red color spectrum at the bottom right. Green to black color bars indicate no significant protein expression level changes, and black to red color bars indicate elevated up-regulation. X-axis top: concentration of the water samples in units of concentration factors. X-axis bottom: treatment process. Y-axis right: list of protein biomarkers color-coded based on their associated stress categories (red: oxidative stress; blue: DNA stress) (b) Summary of integrated molecular toxicity endpoints (TEQgeno and TEQoxi) for DNA and oxidative stress of water samples with high halogen that were treated by chlorination (Cl_2_), chlorine/chloramines (Cl_2_/NH_2_Cl), or GAC with pre- and post-chlorination (Cl_2_/GAC/Cl_2_) from GAC system at different BVs.

For Cl_2_/GAC/Cl_2_ under high halogen levels, unlike cytotoxicity results, we observed no consistent reduction in either oxidative stress or DNA stress across the GAC lifetime, compared to samples treated with Cl_2_/NH_2_Cl, which is inconsistent with the cytotoxicity results. (Figure 5b) The underlying reasons for this inconsistency remain unclear.

Dickenson et al., suggested that Cl_2_/GAC/Cl_2_ indeed increase the calculated genotoxicity of drinking water samples compared to the raw water.(Verdugo et al., 2020) While these results do not align with our previous findings on cytotoxicity, various factors might influence the correlation between different toxicity endpoints. Furthermore, while the cytotoxicity of DBPs is additive, research indicates that the genotoxicity of DBPs in mixtures is less than additive, suggesting an antagonistic interaction whose mechanism is not yet understood.(Lau et al., 2023c) Most studies on the genotoxicity of DBPs have focused on aliphatic DBPs, which constitute less than 50% of DBPs in drinking water; thus, the presence of aromatic DBPs could significantly impact overall toxicity.(Han et al., 2021) However, the challenges in enriching and separating both aliphatic and aromatic DBPs in drinking water, coupled with the lack of detailed knowledge about the genotoxicity ranking of aromatic DBPs and the mechanisms of antagonistic interactions in DBP mixtures, underscore the need for further research to elucidate the chemical and biological mechanisms driving the genotoxicity of drinking water influenced by DBPs.

#### 3.1.2 RT-qPCR assay in human biomarkers

RT-qPCR was utilized to assess the transcriptional changes in four key stress gene biomarkers in human HepG2 cells exposed to 21 drinking water samples at a CF of 100 (Figure S2). Generally, HO1, indicative of oxidative stress, showed higher expression levels compared to KU70 (DNA stress), p53 (apoptosis), and MTF-1 (chemical stress). After 4 hours of exposure, all genes, except for three samples concerning apoptosis and one for DNA damage at BV 16000, were upregulated. HO1 was upregulated in all samples, regardless of the treatment process.

KU70 was upregulated in most samples, except in those treated with Cl_2_/GAC/Cl_2_ at BV 16000. MTF-1 showed increased expression in samples from the GAC system at BV 10 and 16000, whereas no significant changes were observed in samples from BV 6800. p53 was upregulated in samples from the GAC system at BV 10 and 6800; however, two out of four samples were downregulated in GAC influent samples from BV 16000, one with low halide concentration treated with Cl_2_/NH_2_Cl, and another with high halide concentration treated with Cl_2_. It is noteworthy that the relationship between gene expression fold change and sample concentration factor is non-monotonic, with gene expression potentially shifting from upregulation to downregulation within certain concentration ranges (Lan et al., 2018).

Hierarchical clustering analysis revealed three major clusters that define groups of drinking water samples inducing similar gene expression patterns in human cells (Figure 6). The first cluster contained 3 post-GAC samples collected at BV 10, the second cluster included 3 post-GAC samples collected at BV 6800, and the last cluster contained all pre-GAC samples and post-GAC samples collected at BV 16000. Figure 6 demonstrated a clear separation between post-GAC and pre-GAC samples. Additionally, bed volume significantly influenced gene expression, with pronounced differences observed at BV 10 and BV 6800, indicating an alteration in GAC’s effectiveness in reducing treated water toxicity over its lifetime.

**Figure 6:**
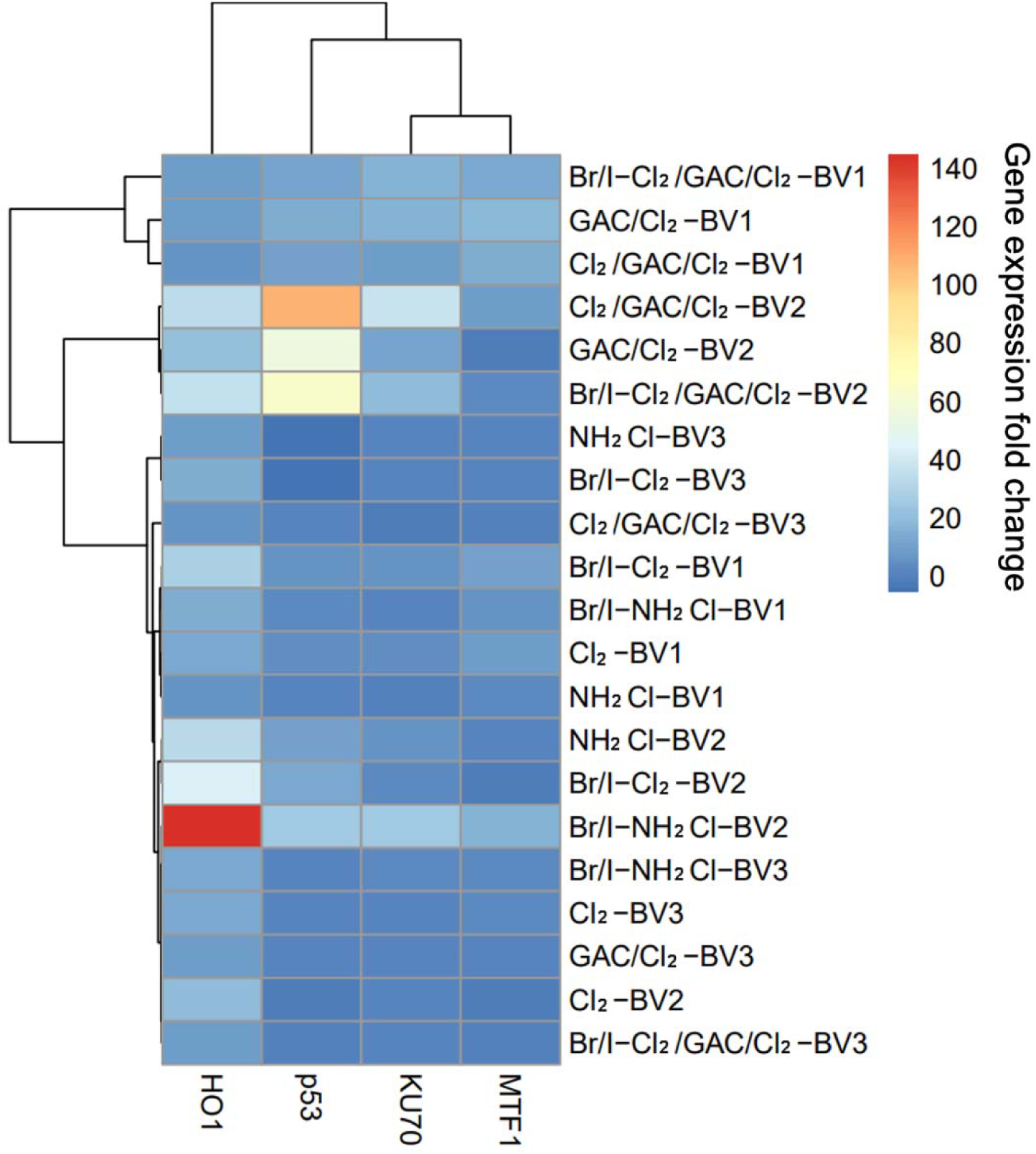
Hierarchical clustering analysis (HCA) diagram based on differential gene expressions of the 4 stress biomarkers in human cell in exposure to drinking water extracts (REF = 100) across different disinfection treatment. X-axis top: cluster root of the biomarkers. X-axis bottom: list of 4 stress biomarkers in human cell. Y-axis left: cluster root of the samples to present three main clusters. Y-axis right: sample names. Br/I represent high halogen condition, while BV 1, 2 and 3 represent bed volume 10, 6800 and 16000, respectively.

### 3.2 Molecular insights into the potential antibiotic resistance induced by drinking water

We performed a GFP-fused E. coli gene ensemble library targeting 6 key antibiotic resistance pathways to assess the resistance induction potential of treated drinking water samples. We employed ARIPI to quantitatively quantify the chemical-induced gene expression level change of a treatment. Although the overall ARIPI values for each sample were below the significant threshold of 1.5, most samples demonstrated significantly elevated ARIPI values compared to the control, except for those treated with both chlorine and chloramine at low halogen levels.

This observation indicates potential antibiotic resistance induction by the samples, highlighting the need for further detailed investigation into the specific role and impact of disinfection processes in promoting antibiotic resistance. (Figure 7)

**Figure 7:**
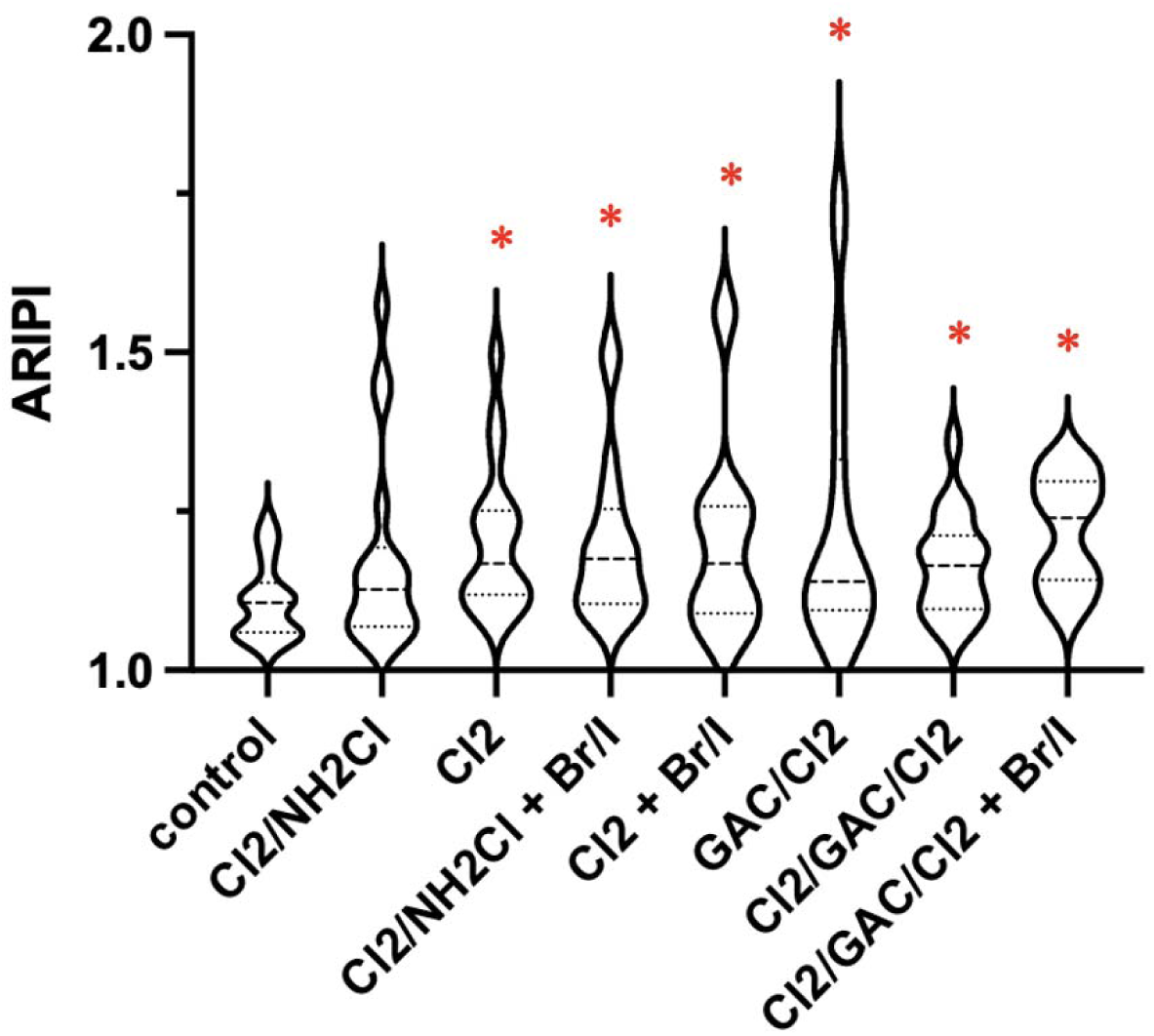
Box plot showing statistics of Antimicrobial Resistance Induction Potential Index (ARIPI) of all 18 genes in the antibiotic resistance induction potential assessment assay in exposure to drinking water samples. Asterisks indicating a significant difference between the tested group and the control.

### 3.3 Insights into the association of detected DBPs with molecular toxicity quantifiers

#### 3.3.1 Correlation analysis between PELI_ORF_ values of yeast biomarkers and detected DBPs

To elucidate the toxicity mechanisms of identified DBPs in disinfected drinking water samples, Spearman correlation was conducted between the concentrations of detected DBPs and toxicity quantifiers obtained from yeast assay. (Figure 8a) For DBPs that were detected below detection limit, half of the detection limit was used based on literature.(Zeng and Mitch, 2016) Among 31 detected DBPs in all the drinking water extracts, 20 of them were significantly corelated with at least 1 biomarker in the yeast library. DBAN, previously shown to impact DNA repair, demonstrated a moderate and significant correlation with RAD27 in our results, a biomarker associated with the base excision repair pathway.(Komaki and Ibuki, 2022)

**Figure 8:**
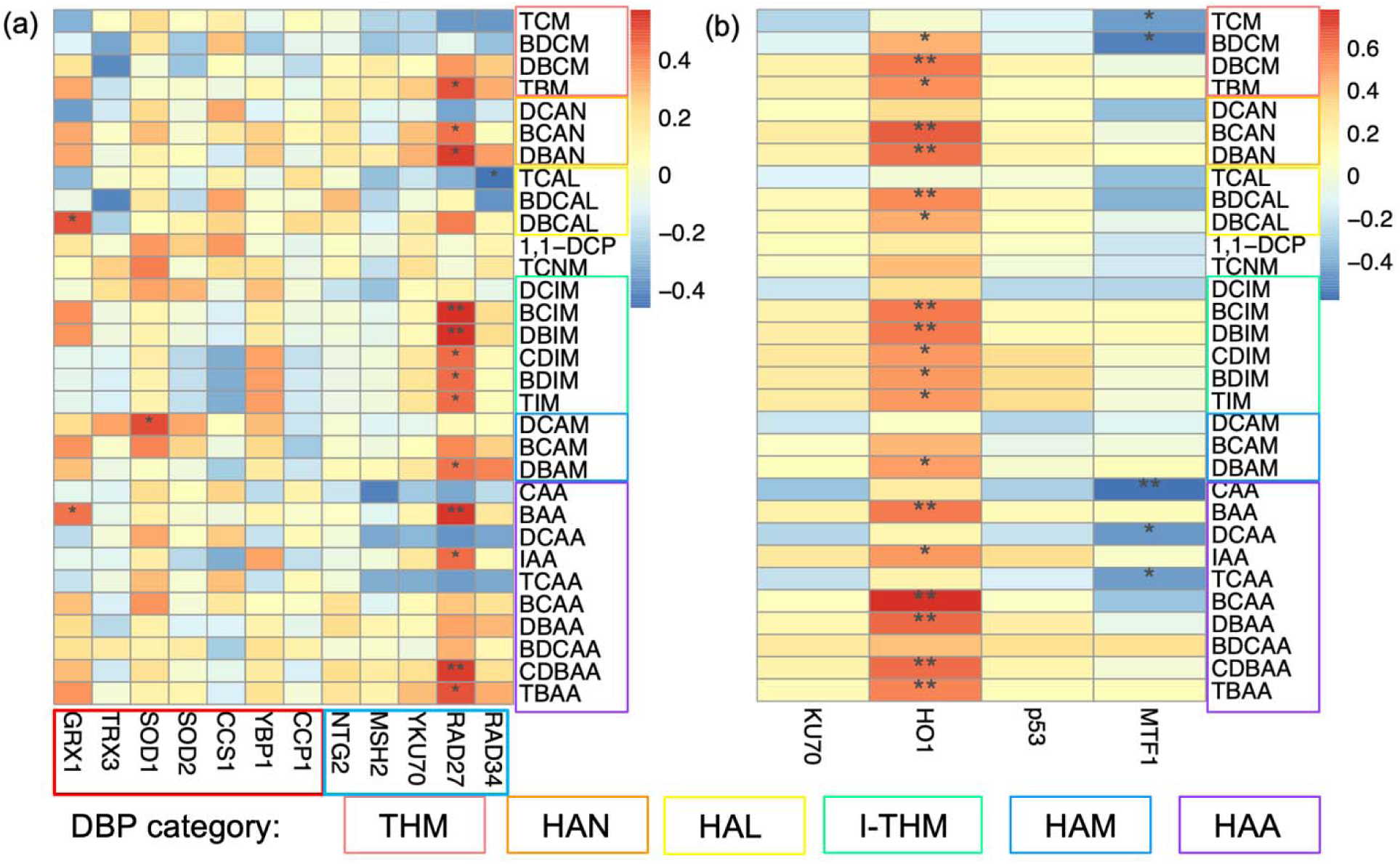
Spearman correlation analysis between (a) DBP concentrations and yeast PELI values for each biomarker for both oxidative stress (red) and DNA stress (blue) and (b) DBP concentrations and molecular toxicity quantifiers of human biomarkers at concentration factor of 100 in drinking water sample extracts. Correlation coefficients at 95% significance level are scaled with color spectrum on the right, zeros are assigned to the non-significant correlations. Asterisks indicating a significant correlation (p<0.05).

Interestingly, DBPs that showed significant correlation with yeast reporters mainly belong some specific DBP categories, namely I-THMs, Haloacetonitriles (HANs) and HAAs (especially the unregulated ones). The Comparative Toxicogenomics Database (CTD) provided further evidence supporting the significance of these DBP categories.(Davis et al., 2023) For instance, DBAN shares a comparable set of interacting proteins with BCAN, another DBP that also showed significant correlation with RAD27. Both DBAN and BCAN are nitrogenous DBPs (N-DBPs) and belong to the HANs group, which is known to exhibit higher genotoxicity than C-DBPs, with HANs generally being the most toxic among N-DBPs. (Lan et al., 2018; Plewa et al., 2008)

Overall, these DBPs were proved to be more genotoxic than the regulated Cl-DBPs in the literature, indicating that I-DBPs may be the main toxic drivers in DNA stress related pathway.(Plewa et al., 2008; Plewa et al., 2017)

#### 3.3.2 Correlation analysis between molecular toxicity quantifiers of human biomarkers and detected DBPs

Spearman correlation was also conducted between the concentrations of detected DBPs and toxicity quantifiers obtained from human biomarkers to elucidate the toxicity mechanisms of identified DBPs in disinfected drinking water samples. (Figure 8b) Similar to the correlation observed between stress responses in yeast and DBP concentrations, the correlation between human biomarkers and DBPs highlighted similar DBP categories. Our result also suggested that all the detected I-THMs were significantly correlated with HO1, while the CTD database suggested that most of the THMs shared similar toxicity pathways including biological oxidation, apoptosis, and pathways in cancer; and the related phenotype including the response to oxidative stress, which is similar with our result. (Figure 8b).

Unlike the results in the yeast assay, HO1 is the only biomarker found to be positively significantly corelated with detected DBPs. Similar with the correlation results in yeast assay, I-THMs and HANs were two of the major categories of DBPs that were found to be significantly correlated with HO1, indicating that these two DBP classes may the main toxic drivers of oxidative stress in human cells. Additionally, some regulated DBPs, such as DBAA and BAA, were also significantly correlated with HO1, emphasizing the need for continued monitoring and regulation of these compounds in treated drinking water.

The consistent findings from both yeast and human cell assays highlight significant concerns regarding multiple DBP categories, particularly I-THMs and HANs. Furthermore, the significant correlation of regulated DBPs like DBAA and BAA with human toxicity markers suggests that regulated DBPs can also contribute to adverse health effects, highlighting the importance of strict regulation and monitoring.

### 3.4 Application of maximum cumulative ratio (MCR) approach identifies cumulative risk driver

Due to the growing concern and uncertain on the potential effect of the mixture of DBPs in drinking water system, a tiered approach incorporating the mixture characterization ratio (MCR) concept was employed for the risk assessment of treated drinking water. The risk assessment results, including hazard indices (HIs), hazard quotients (HQs), and MCRs for each sample, are summarized in Table 1. (Vallotton and Price, 2016)

**Table 1:** Hazard Index (HI), Maximum Hazard Quotient (HQMAX), Maximum Cumulative Ratio (MCR), and top five high-priority DBPs (highest HQ values) in the 21 disinfected water samples collected from the GAC pilot system.

| **Bed volume** | **Halide concentration** | **Disinfection process** | **HI** | **HQ max** | **MCR** | **Top 5 High-Priority DBPs** |
| --- | --- | --- | --- | --- | --- | --- |
| **Bed volume 10** | Without Br/I | NH<sub>2</sub>Cl/Cl<sub>2</sub> | 0.327 | 0.124 | 2.646 | THM5, HAA4, IAA, DBAN, BCAM |
|  | Without Br/I | Cl<sub>2</sub> | 0.691 | 0.335 | 2.062 | THM5, HAA4, IAA, DBAN, DBCAL |
|  | With Br/I | NH<sub>2</sub>Cl/Cl<sub>2</sub> | 0.778 | 0.274 | 2.843 | THM5, HAA4, DBAN, DBAM, BCAM |
|  | With Br/I | Cl<sub>2</sub> | 1.365 | 0.482 | 2.830 | THM5, HAA4, DBAN, DBCAL, DBAM |
|  | Without Br/I | GAC/Cl<sub>2</sub> | 0.093 | 0.021 | 4.400 | THM5, HAA4, IAA, DBAN, DBCAL |
|  | Without Br/I | Cl<sub>2</sub>/GAC/Cl<sub>2</sub> | 0.124 | 0.041 | 3.001 | THM5, HAA4, IAA, DBAN, DBCAL |
|  | With Br/I | Cl<sub>2</sub>/GAC/Cl<sub>2</sub> | 0.391 | 0.175 | 2.234 | THM5, IAA, HAA4, DBAN, DBCAL |
| **Bed volume 6800** | Without Br/I | NH<sub>2</sub>Cl/Cl<sub>2</sub> | 0.793 | 0.302 | 2.623 | THM5, HAA4, BCAM, IAA, DBAN |
|  | Without Br/I | Cl<sub>2</sub> | 1.974 | 0.900 | 2.193 | THM5, HAA4, IAA, BDCAL, BCAN |
|  | With Br/I | NH<sub>2</sub>Cl/Cl<sub>2</sub> | 1.379 | 0.294 | 4.695 | THM5, DBAN, HAA4, DBAM, IAA |
|  | With Br/I | Cl<sub>2</sub> | 3.497 | 0.814 | 4.296 | DBCAL, THM5, DBAN, HAA4, BDCAL |
|  | Without Br/I | Cl<sub>2</sub>/GAC | 0.749 | 0.337 | 2.219 | THM5, HAA4, BDCAL, IAA, BCAN |
|  | Without Br/I | Cl<sub>2</sub>/GAC/Cl<sub>2</sub> | 0.635 | 0.294 | 2.162 | THM5, HAA4, IAA, DBAN, BDCAL |
|  | With Br/I | Cl<sub>2</sub>/GAC/Cl<sub>2</sub> | 1.279 | 0.375 | 3.411 | THM5, DBCAL, DBAN, HAA4, DBAM |
| **Bed volume 16000** | Without Br/I | NH<sub>2</sub>Cl/Cl<sub>2</sub> | 0.464 | 0.181 | 2.561 | THM5, HAA4, BCAM, IAA, DBAN |
|  | Without Br/I | Cl<sub>2</sub> | 1.020 | 0.451 | 2.260 | THM5, HAA4, IAA, DBAN, DBCAL |
|  | With Br/I | NH<sub>2</sub>Cl/Cl<sub>2</sub> | 0.653 | 0.194 | 3.372 | THM5, HAA4, DBAM, DBAN, DBCAL |
|  | With Br/I | Cl<sub>2</sub> | 1.288 | 0.416 | 3.095 | THM5, HAA4, DBAN, DBCAL, DBAM |
|  | Without Br/I | Cl<sub>2</sub>/GAC | 0.608 | 0.271 | 2.243 | THM5, HAA4, IAA, DBAN, DBCAL |
|  | Without Br/I | Cl<sub>2</sub>/GAC/Cl<sub>2</sub> | 0.592 | 0.286 | 2.069 | THM5, HAA4, IAA, DBAN, DBCAL |
|  | With Br/I | Cl<sub>2</sub>/GAC/Cl<sub>2</sub> | 0.621 | 0.274 | 2.271 | THM5, HAA4, DBAN, DBAM, DBCAL |

For all low halogen concentration samples disinfected with chlorine and chloramine, the HI and HQ for all three samples were under risk group II, indicating low or no concern for the potential cumulative adverse health effects in these samples. For the low halogen concentration samples disinfected by chlorine only, sample taken at BV 10 was under risk group II, while samples taken at BV 6800 and 16000 were under risk group IIIB, indicating that the identified potential risks offered by the samples may be driven by multi DBPs. For the pre-GAC samples with high halogen concentration, samples treated with chlorine only were under risk group IIIB, while the only sample treated with both chlorine and chloramine under BV 6800 was under risk group IIIB, rest of them were under risk group II. For the post-GAC samples, HI of nearly all effluent samples were <1 (risk group II), expect the spiked samples treated with Cl_2_/GAC/Cl_2_ under BV 6800 (risk group IIIB).

Moreover, to identify the major toxicity drivers in drinking water treated with different disinfection process, individual DBPs were ranked based on their HQ, and the results were listed in the right column of Table 1. Regulated DBP classes (TTHM and HAA5) were both the top two predominant contaminants in all samples. It is worth noting that the risk of TTHM and HAA5 were calculated based on the total concentration of them instead of individual DBP species, which would potentially result in a higher HQ value compared to the unregulated ones. In addition, although regulated DBPs are less toxic than the unregulated ones, their concentration in disinfected water are generally higher than the unregulated ones, similar trend also identified in our results, which also explain the results that the higher health risk of regulated DBPs than unregulated ones that we concluded. (Allen et al., 2021)

Furthermore, among all the unregulated DBPs target in this study, IAA, DBAN, DBCAL, BCAN, BCAM were highly counted emerging DBPs in the top 5 high-priority list. IAA was I-DBP, DBAN, DBCAL, BCAN, BCAM were Br-DBP or N-DBP, restating the previously reported concerns regarding these DBP classes.(Hanigan et al., 2017; Richardson et al., 2007) However, almost all unregulated DBPs had detected concentration less than one-tenth of their human-health benchmarks (HQ < 0.1), with 2 exception, DBAN had detected concentration greater than one-tenth of their human-health benchmarks in high halogen samples collected from BV 6800(0.1 < HQ ≤ 1).

## 4 Conclusion

This study demonstrated how different treatment and disinfection processes impact the toxicity of drinking water, illustrating the complex interplay between GAC treatment, DBP formation, and the subsequent toxicity of treated drinking water. Our results suggested that at low halogen conditions, GAC/Cl_2_ generally reduces both genotoxicity and oxidative stress more effectively than Cl_2_/NH_2_Cl. However, Cl_2_/GAC/Cl_2_ does not consistently lower toxicity and, under high halogen conditions, fails to mitigate genotoxicity and oxidative stress compared to GAC/Cl2. Correlation analysis indicated that the genotoxic effects and oxidative stress effect in treated drinking water are significantly influenced by unregulated DBPs (especially I-THMs and HANs). This highlights the complex dynamics of water treatment and emphasizes the critical need to address the impact of these unregulated and unknown DBPs on water safety. We also observed elevated levels in samples with various disinfection and treatment processes, indicating a potential risk for antibiotic resistance induction, warranting further investigation into the mechanisms by which disinfection processes may contribute to this emerging public health concern. Moreover, future studies might be necessary to unravel the mechanisms of DBP mixture induced genotoxicity and oxidative stress to ensure the ongoing challenges in the safety of treated drinking water, and to reduce the toxicological impacts of treated drinking water while meeting regulatory standards.

## Acknowledgement

This work was supported by funding from the Water Research Foundation (WRF 5140).

