## Supplemental Information for "Impacts of Disinfection Methods in a Granular Activated Carbon (GAC) Treatment System on Disinfected Drinking Water Toxicity and Antibiotic Resistance Induction Potential"

List of Table:

### Text S1: GAC pilot system

Three parallel 6.4-cm diameter granular activated carbon (GAC) columns were operated to evaluate different conditions (Figure 1) including Column 1: GAC treatment without bromide (Br-) and iodide (I-) spiking or pre-chlorination, Column 2: GAC treatment without Br- and I- spiking but with pre-chlorination, and Column 3: GAC treatment with Br- and I- spiking and pre-chlorination; each column contained 81 cm of Filtrasorb 400-M (0.55–0.75 effective grain size; Calgon Carbon Corporation), resulting in a 2.6 L empty bed volume and operated at a hydraulic loading rate of 1.36 mm/s (2.0 gpm/ft2) to achieve a 10-minute empty bed contact time (EBCT); bromide and iodide were added using a peristaltic pump from a stock solution, with measured Br- concentrations ranging from 190-260 µg/L and estimated I- concentrations ranging from 57-78 µg/L based on the concentration ratio in the stock solution. For Columns 2 and 3, chlorine was introduced to attain a residual concentration of 0.1 mg/L as Cl2 after a 2-hour contact time which was achieved by passing the water through a reactor upstream of the GAC columns.

### Text S2: Sample preparation and extraction

10 L of samples from GAC system at each sampling location were treated with chlorine only or chlorine/chloramines (dose ≈ 2 mg/L as Cl2). After 3 days of contact, the residual (≥ 1 mg/L as Cl2) was quenched using sodium thiosulfate, and the quenched samples were acidified to ~pH 3.7 by sulfuric acid prior to the solid-phase extraction (SPE) process.

Custom-made solid-phase extraction (SPE) cartridges were prepared, with each cartridge containing 2.5 g of Sepra ZTL (Phenomenex), a surface-modified styrene divinylbenzene sorbent similar to Strata-X. The 10-L quenched and acidified water samples were divided into two equal 5-L aliquots, and each aliquot was extracted using one SPE cartridge. Prior to sample loading, the cartridges were pre-conditioned with 50 mL of methanol and then equilibrated with 50 mL of deionized (DI) water containing 0.01% (v/v) sulfuric acid. The water samples were loaded onto the SPE cartridges using a vacuum manifold at a flow rate of approximately 11 mL/min. Subsequently, the cartridges were dried under vacuum overnight. The following day, each cartridge was eluted with 40 mL of distilled methanol, and the eluates from the two cartridges for each 10-L water sample were combined and concentrated to less than 10 mL using roto evaporation. The methanol was further concentrated to less than 5 mL using a gentle stream of nitrogen. To perform solvent exchange, 200 μL of dimethyl sulfoxide (DMSO) was added to the methanol, and the DMSO-methanol mixture was concentrated using nitrogen blowdown. The final volume of each DMSO extract was measured to determine the achieved concentration factor of the extractions.

### Text S3: Gene expression profiling data processing and quantitative molecular toxicity endpoints derivation

Temporal raw data of optical density (OD) and green fluorescent protein (GFP) signals are first corrected by subtracting the OD and GFP signals of medium controls and promoterless bacteria controls, both with and without sample exposure, respectively. The induction factor (I), representing the change in gene expression for a given gene at each time point due to exposure to effluent water samples compared to an untreated control, is calculated as I = Pe/Pc, where Pe = (GFP/OD)_experiment_, which is the normalized gene expression GFP level in the experimental condition with sample exposure, and Pc = (GFP/OD)_control_, which is the normalized gene expression GFP level in the control condition without any sample exposure.

To quantify the changes in gene expression levels induced by chemical treatment, we proposed the ARIPI as a molecular toxicity quantifier. The cumulative change in altered gene expression over the 2-hour exposure period was calculated as follows:

$${ARIPI}_{gene i}=\frac{\int_{t=0}^{t} e^{|\ln I|}dt}{exposure time}$$

    Where, t is the exposure time.

### Text S4: Maximum Cumulative Ratio (MCR) approach

Maximum cumulative ratio (MCR), defined as the cumulative health risks of a mixture divided by the maximum health risk of a single component, was employed to identify the health risks and prioritize contaminants in the disinfected water samples. Health risks of individual chemicals and the mixture were expressed as hazard quotients (HQs) and hazard indexes (HIs), respectively. HQs of individual contaminants were derived by comparing the chemical concentrations detected in the environment to their permitted doses. In this study, for total trihalomethanes (TTHMs) and the five haloacetic acids (HAA5), which are regulated in drinking water by U.S. EPA, their permitted doses were selected as regulatory maximum contaminant level (MCL). For other non-regulated chemicals, the LC50 data from a Chinese hamster ovary (CHO) cell chronic cytotoxicity assay reported in literature were used as MCLs. HIs were calculated by summing up all available HQs in the mixture based on the assumption of concentration additoin69. In environmental samples where chemical concentrations are low, concentration addition has been widely accepted as a conservative model due to the rare synergism. To untangle the major risk drivers in the collected samples, MCR was calculated as:

MCR = $\frac{HI}{{HQ}_{max}}$

    where, HQMAX is the maximum HQ of multiple components in the mixture.

The larger the MCR (>2), it is more likely that the health risks of the mixture cannot be simply attributed to one or two major components. Therefore, further investigation is required since a chemical-by-chemical regulation approach would underestimate the overall toxicity of the mixture. Based on their different HQ, HI and MCR values, samples can be categorized into four risk groups as previously described.

### Table S1: List of selected protein biomarkers from the yeast stress response library in this study, and their corresponding stress and toxicity pathway information.(Rahman et al., 2022)

| **Stress** | **Function** | **Pathway** | **Key Biomarker** |
| --- | --- | --- | --- |
| DNA stress | DNA repair | Base excision repair (BER) | NTG2, RAD27 |
|  |  | Nucleotide excision repair (NER) | RAD34 |
|  |  | Mismatch repair (MMR) | MSH2 |
|  |  | Non-homologous end joining (NHEJ) | YKU70 |
| Oxidative stress | Defense system | Superoxide dismutase (SOD) | CCS1, SOD1, SOD2 |
|  |  | Cytochrome C (CYC) related | CCP1 |
|  |  | Thioredoxin | TRX3 |
|  |  | Glutaredoxin | GRX1 |
|  | Sensor/regulator | Yap1p regulation | YBP1 |

### Table S2: List of genes, related pathways and mechanism information in the *E.coli* antibiotic resistance library

| Mechanism | Gene selected | Functions |
| --- | --- | --- |
| Outer membrane permeability | *rfaZ, rcsA, slp, ompN, rob* | Lipid-mediated related, porin related, PBP related, Porin-mediated antibiotic permeability |
| Efflux pump | *entS, oppB, hisJ, gltJ, cusC,* | Major facilitator superfamily, ATP-binding cassette, Resistance-nodulation-cell division resistance family |
| Drug inactivation | *cysD, yncA* | Beta-lactam enzyme genes |
| Targets alternation | *gyrB, sulA* | Quinolones targets, sulfonamides targets |
| SOS response | *yedP, rpoS* | SOS response |
| Detoxification | *katE, osmC* | oxidative stress |

(a)

| Bed volume | Halide condition | Disinfection process | PELIgeno 1.5 | PELIoxi 1.5 |
| --- | --- | --- | --- | --- |
| Bed volume 10 | Without Br/I | Cl2/NH2Cl | 58.61 | 7.39 |
|  |  | Cl2 | 53.83 | 9.70 |
|  | With Br/I | Cl2/NH2Cl | 50.93 | 12.85 |
|  |  | Cl2 | 38.46 | 6.07 |
| Bed volume 6800 | Without Br/I | Cl2/NH2Cl | 9.79 | 75.16 |
|  |  | Cl2 | 15.31 | 51.76 |
|  | With Br/I | Cl2/NH2Cl | 51.28 | 64.56 |
|  |  | Cl2 | 18.62 | 59.02 |
| Bed volume 16000 | Without Br/I | Cl2/NH2Cl | 67.92 | 33.11 |
|  |  | Cl2 | 70.79 | 72.44 |
|  | With Br/I | Cl2/NH2Cl | 75.16 | 28.18 |
|  |  | Cl2 | 95.50 | 35.40 |

(b)

| Bed volume | Halide condition | Treatment process | PELIgeno 1.5 | PELIoxi 1.5 |
| --- | --- | --- | --- | --- |
| Bed volume 10 | Without Br/I | Cl2/NH2Cl | 58.61 | 7.39 |
|  |  | Cl2 | 53.83 | 9.70 |
|  |  | GAC/Cl2 | 66.37 | 11.22 |
| Bed volume 6800 |  | Cl2/NH2Cl | 9.79 | 75.16 |
|  |  | Cl2 | 15.31 | 51.76 |
|  |  | GAC/Cl2 | 28.97 | 86.10 |
| Bed volume 16000 |  | Cl2/NH2Cl | 67.92 | 33.11 |
|  |  | Cl2 | 70.79 | 72.44 |
|  |  | GAC/Cl2 | 149.28 | 14.93 |

(c)

| Bed volume | Halide condition | GAC treatment process | PELIgeno 1.5 | PELIoxi 1.5 |
| --- | --- | --- | --- | --- |
| Bed volume 10 | Without Br/I | Cl2 | 53.83 | 9.70 |
|  |  | GAC/Cl2 | 66.37 | 11.22 |
|  |  | Cl2/ GAC/Cl2 | 77.80 | 36.06 |
| Bed volume 6800 |  | Cl2 | 15.31 | 51.76 |
|  |  | GAC/Cl2 | 28.97 | 86.10 |
|  |  | Cl2/ GAC/Cl2 | 138.99 | 233.36 |
| Bed volume 16000 |  | Cl2 | 70.79 | 72.44 |
|  |  | GAC/Cl2 | 149.28 | 14.93 |
|  |  | Cl2/ GAC/Cl2 | 123.88 | 16.07 |

(d)

| Bed volume | Halide condition | Treatment process | PELIgeno 1.5 | PELIoxi 1.5 |
| --- | --- | --- | --- | --- |
| Bed volume 10 | With Br/I | Cl2/NH2Cl | 50.93 | 12.85 |
|  |  | Cl2 | 38.46 | 6.07 |
|  |  | Cl2/ GAC/Cl2 | 97.70 | 64.56 |
| Bed volume 6800 |  | Cl2/NH2Cl | 51.28 | 64.56 |
|  |  | Cl2 | 18.62 | 59.02 |
|  |  | Cl2/ GAC/Cl2 | 4.69 | 6.02 |
| Bed volume 16000 |  | Cl2/NH2Cl | 75.16 | 28.18 |
|  |  | Cl2 | 95.50 | 35.40 |
|  |  | Cl2/ GAC/Cl2 | 6.28 | 73.45 |

Table S3: Summary of integrated molecular toxicity endpoints (PELIgeno1.5 and PELIoxi1.5) for DNA and oxidative stress of water samples treated with different disinfection processes. (a) GAC influents treated with free chlorine only (Cl2) and chlorine/chloramines (Cl2/NH2Cl) from GAC system at different bed volumes, (b) water samples without Br– and I– spiking that were treated by chlorination (Cl2), GAC with post-chlorination (GAC/Cl2), or chlorination/chloramination (Cl2/NH2Cl) from GAC system at different bed volumes, (c) water samples without Br– and I– spiking that were treated by chlorination (Cl2), GAC with post-chlorination (GAC/Cl2), or GAC with pre- and post-chlorination (Cl2/GAC/Cl2) from GAC system at different bed volumes, and (d) water samples with Br– and I– spiking that were treated by chlorination (Cl2), chlorine/chloramines (Cl2/NH2Cl), or GAC with pre- and post-chlorination (Cl2/GAC/Cl2) from GAC system at different bed volumes

### Table S4: Risk groups based on hazard quotient (HQ), hazard index (HI) and maximum accumulative ratio (MCR) values

| **Group** | **Cumulated mixture risk** | **Individual chemical risk** | **MCR** | **Implications** |
| --- | --- | --- | --- | --- |
| I | HI > 1 | HQ_MAX_ > 1 |  | The mixture presents a  potential risk already based  on individual components |
| II | HI < 1 | HQ_MAX_ < 1 |  | The assessment does not  identify a health risk concern |
| IIIA | HI > 1 | HQ_MAX_ < 1 | MCR < 2 | The majority of the health risk  caused by the mixture is  driven by one contaminant |
| IIIB | HI > 1 | HQ_MAX_ < 1 | MCR > 2 | The potential health risk is driven by multiple contaminants in a specific sample |

### Table S5: Toxicity endpoints of detected DBPs used for MCR approach in this study

| **DBP class** | **DBP** | molecular weight | LC50 umol/L | LC50 ug/L | MCL ug/L |
| --- | --- | --- | --- | --- | --- |
| THM | TCM | 119.37 | 9620 | 1148339.4 | 80 |
| THM | BDCM | 163.83 | 11500 | 1884045 | 80 |
| THM | DBCM | 208.28 | 5360 | 1116380.8 | 80 |
| THM | TBM | 252.73 | 3960 | 1000810.8 | 80 |
| HAN | TCAN | 144.38 | 160 | 23100.8 | 1040 |
| HAN | DCAN | 109.94 | 57.3 | 6299.562 | 280 |
| HAN | BCAN | 154.39 | 8.46 | 1306.1394 | **60** |
| HAN | DBAN | 198.84 | 2.85 | 566.694 | 30 |
| HAL | TCAL | 147.38 | 1163 | 171402.94 | 7710 |
| HAL | BDCAL | 191.83 | 20.35 | 3903.7405 | 180 |
| HAL | DBCAL | 236.29 | 5.15 | 1216.8935 | 50 |
| HAL | TBAL | 280.74 | 3.58 | 1005.0492 | 50 |
| HK | 1,1-DCP |  | NA^a^ |  |  |
| HK | 1,1,1-TCP |  | NA |  |  |
| HNM | TCNM | 164.37 | 536 | 88102.32 | 3960 |
| I-THM | DCIM | 210.82 | 4130 | 870686.6 | 39160 |
| I-THM | BCIM | 255.28 | 2420 | 617777.6 | 27790 |
| I-THM | DBIM | 299.73 | 1910 | 572484.3 | 25750 |
| I-THM | CDIM | 302.28 | 2410 | 728494.8 | 32770 |
| I-THM | BDIM | 346.73 | 1400 | 485422 | 21830 |
| I-THM | TIM | 393.73 | 66 | 25986.18 | 1170 |
| HAM | DCAM | 127.95 | 1920 | 245664 | 11050 |
| HAM | BCAM | 216.86 | 17.1 | 3708.306 | 130 |
| HAM | TCAM | 162.39 | 2050 | 332899.5 | 14970 |
| HAM | DBAM | 216.86 | 12.2 | 2645.692 | 120 |
| HAA | CAA | 94.5 | 810 | 76545 | 60 |
| HAA | BAA | 138.95 | 9.6 | 1333.92 | 60 |
| HAA | DCAA | 128.94 | 7300 | 941262 | 60 |
| HAA | IAA | 185.95 | 2.95 | 548.5525 | 20 |
| HAA | TCAA | 163.38 | 2400 | 392112 | 60 |
| HAA | BCAA | 173.39 | 778 | 134897.42 | 6070 |
| HAA | DBAA | 217.84 | 590 | 128525.6 | 60 |
| HAA | BDCAA | 207.83 | 685 | 142363.55 | 6400 |
| HAA | CDBAA | 252.29 | 202 | 50962.58 | 2290 |
| HAA | TBAA | 296.74 | 85 | 25222.9 | 1130 |

a: NA indicating not detected in all the samples in this study

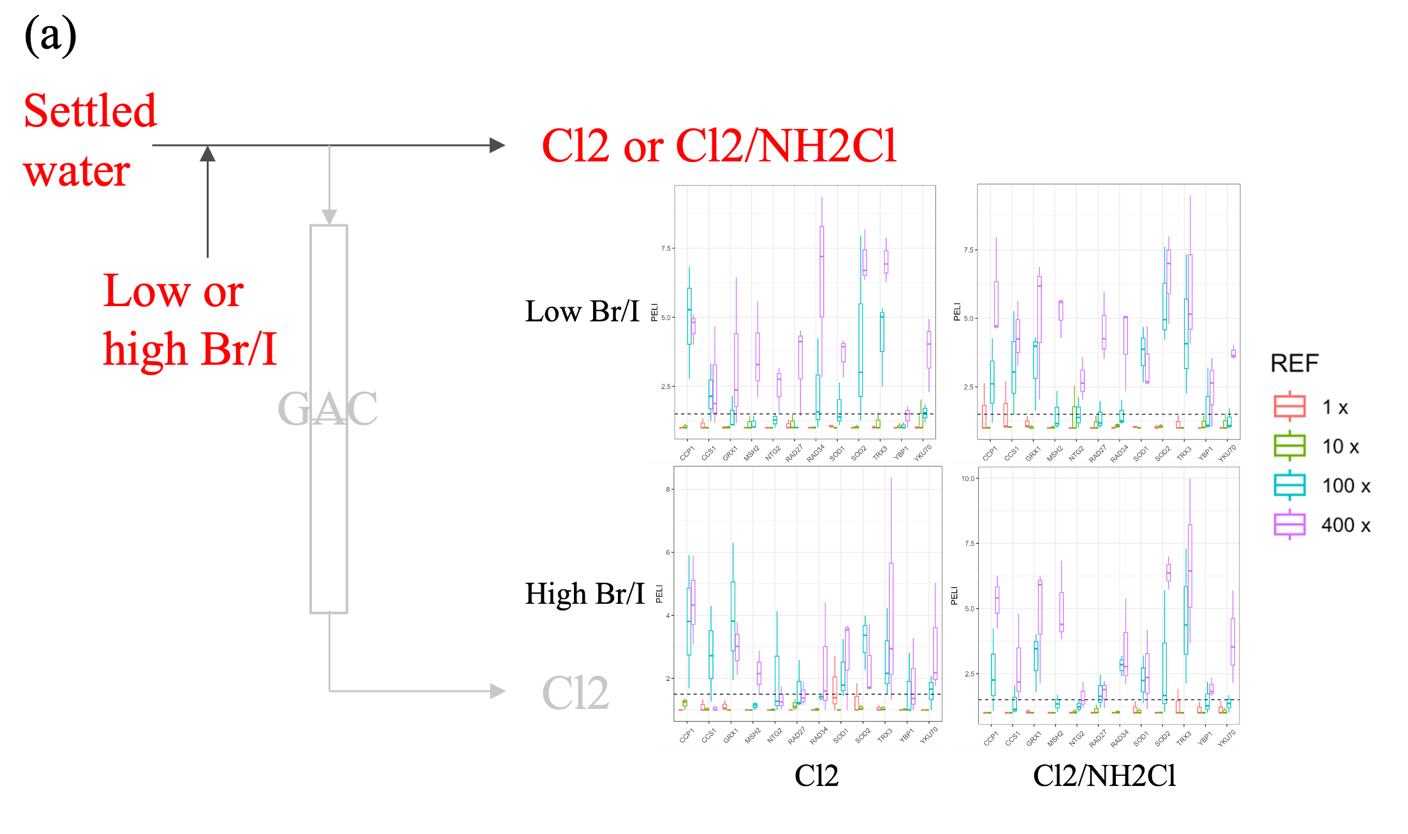

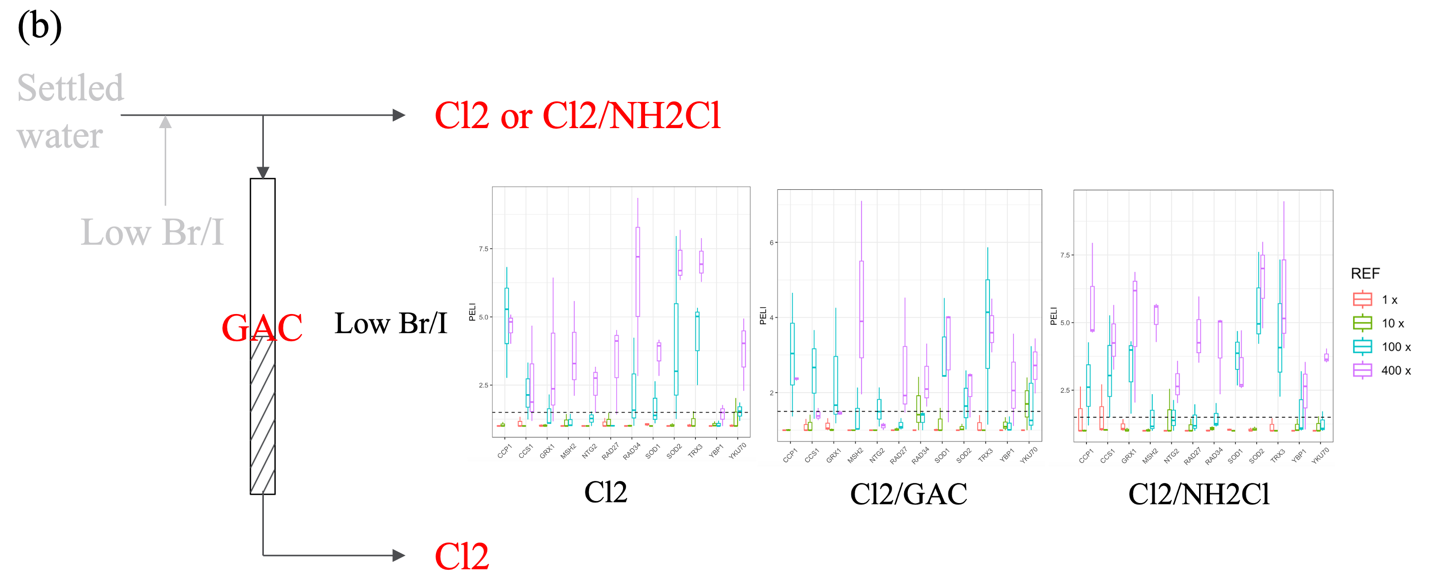

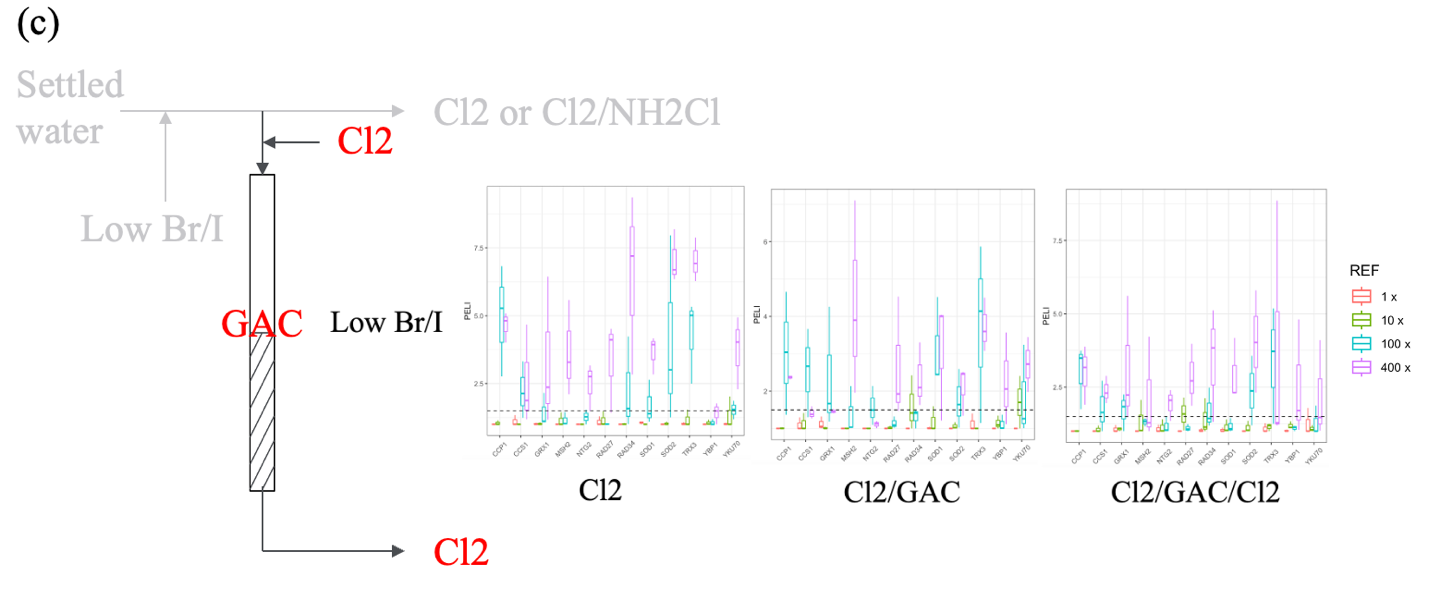

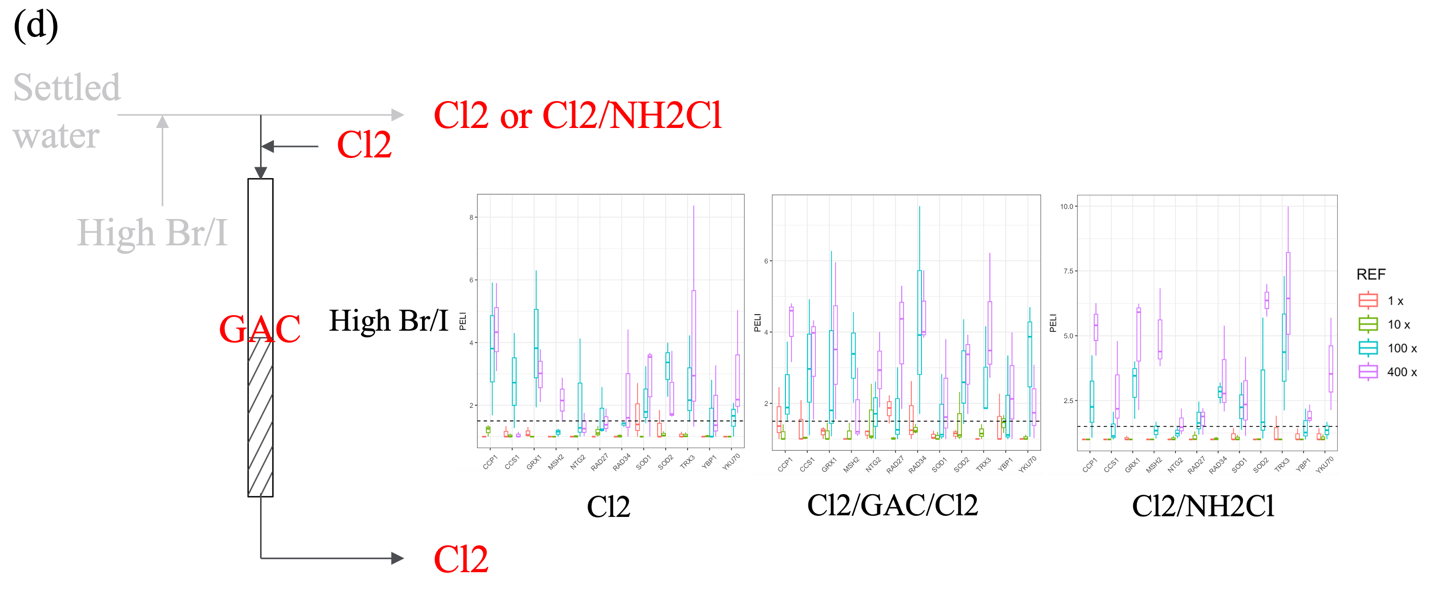

Figure S1: Box plot showing the gene expression of GFP-fused yeast reporter library targeting 12 biomarkers involved in DNA stress and oxidative stress after exposure under drinking water with different treatment process (a) GAC influents treated with free chlorine only (Cl2) and chlorine/chloramines (Cl2/NH2Cl) from GAC system, (b) water samples without Br– and I– spiking that were treated by chlorination (Cl2), GAC with post-chlorination (GAC+Cl2), or chlorination/chloramination (Cl2/NH2Cl) from GAC system, (c) water samples without Br– and I– spiking that were treated by chlorination (Cl2), GAC with post-chlorination (GAC+Cl2), or GAC with pre- and post-chlorination (Cl2+GAC+Cl2) from GAC system across four concentrations, and (d) water samples with Br– and I– spiking that were treated by chlorination (Cl2), chlorine/chloramines (Cl2/NH2Cl), or GAC with pre- and post-chlorination (Cl2/GAC/Cl2) from GAC system

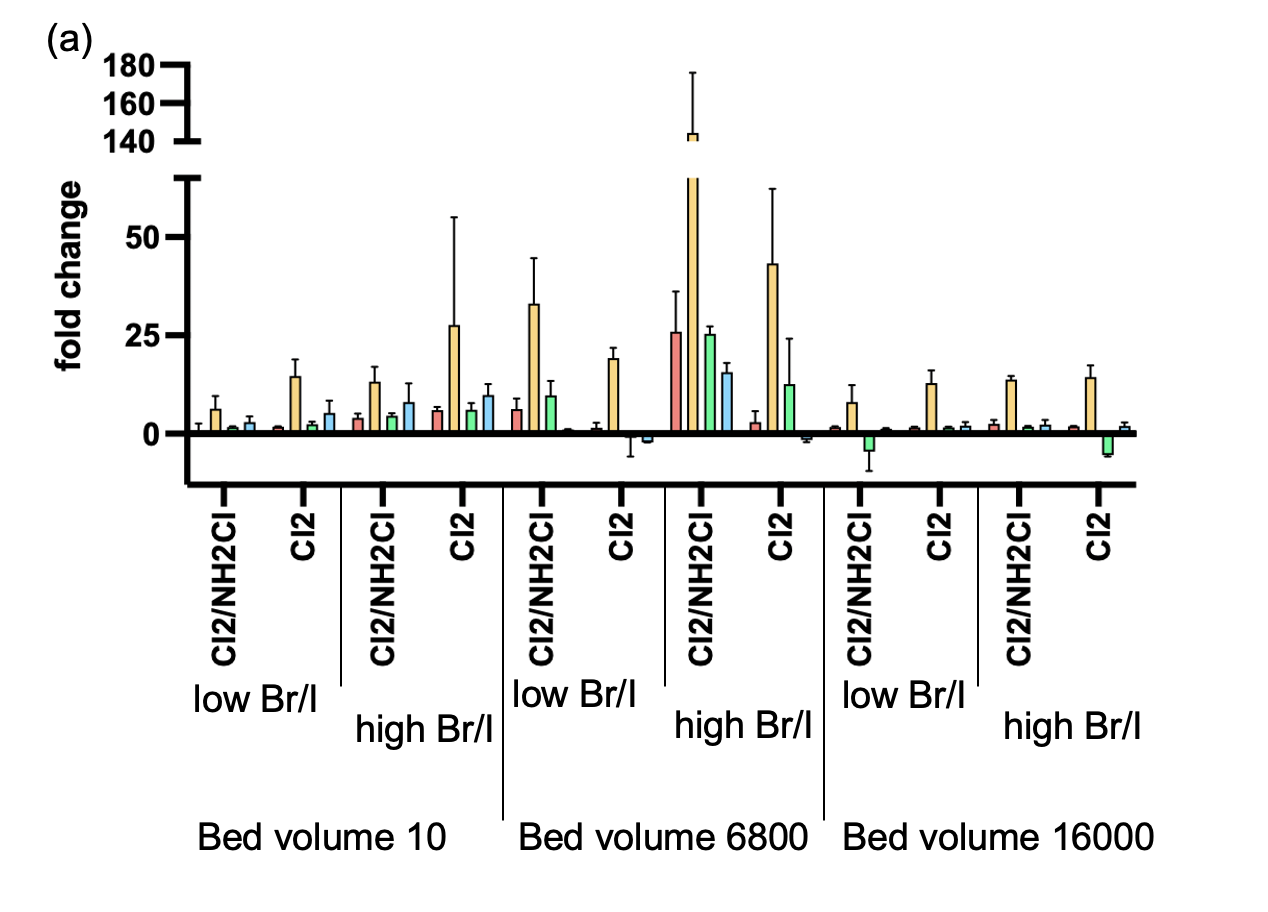

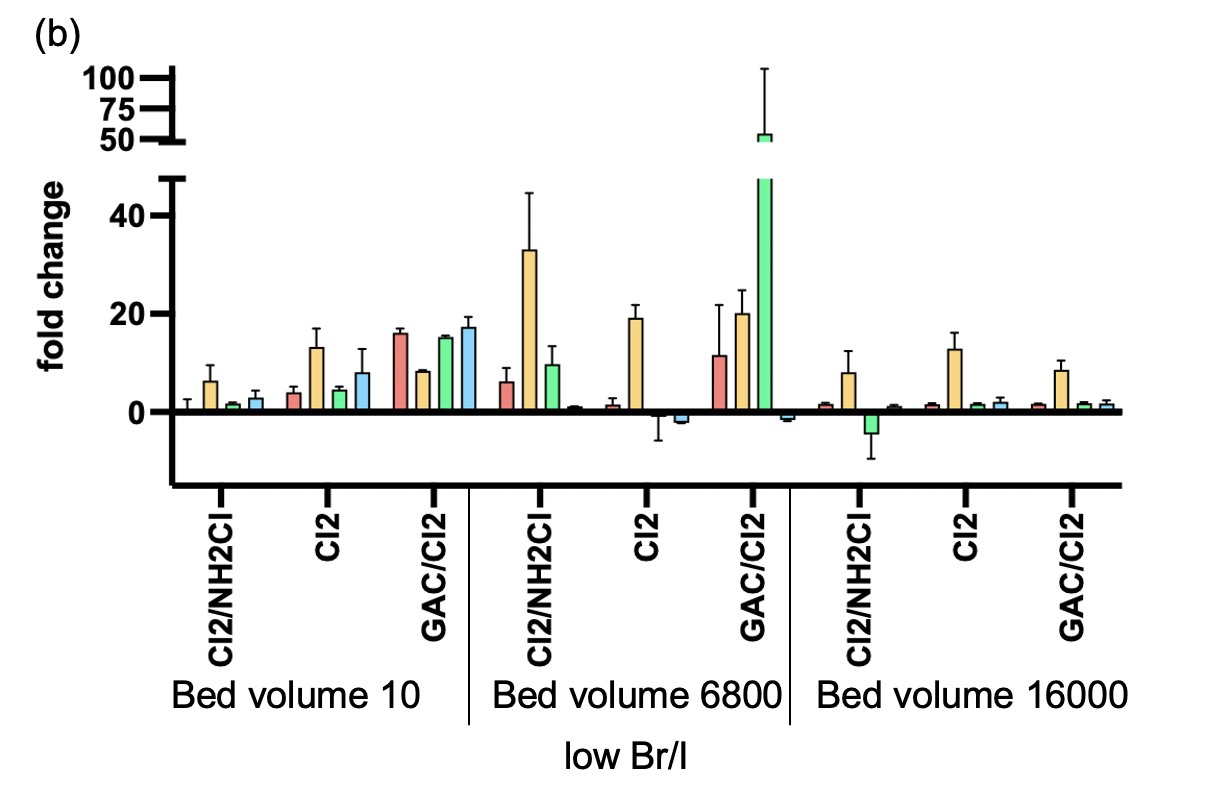

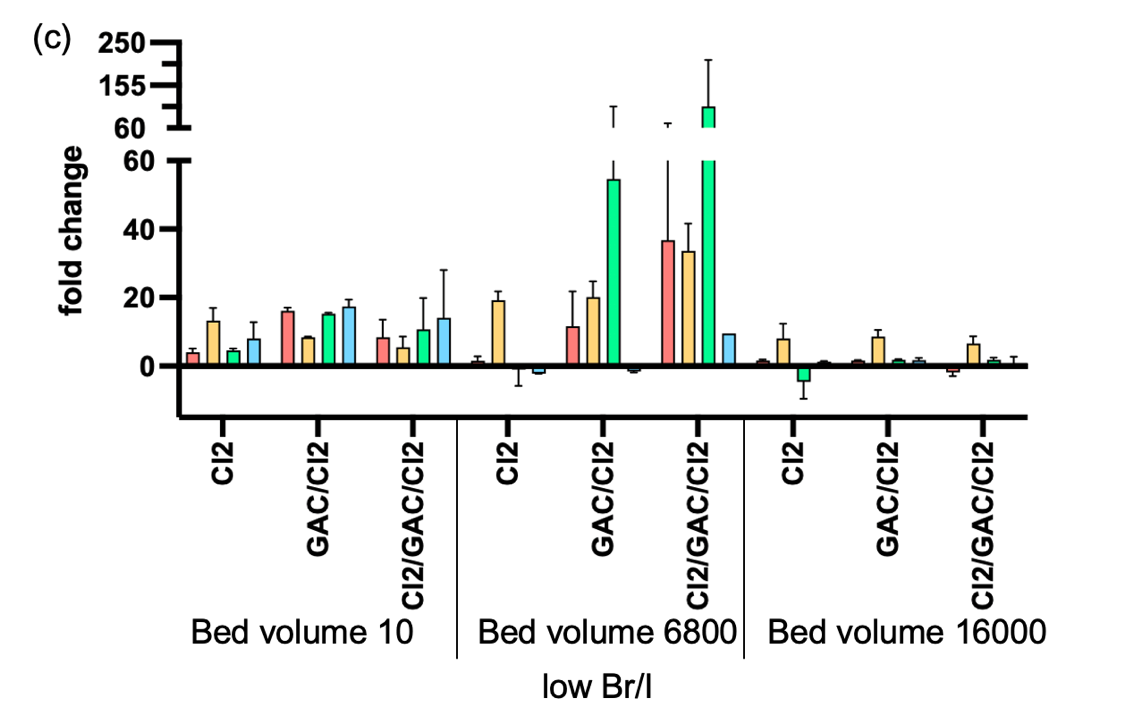

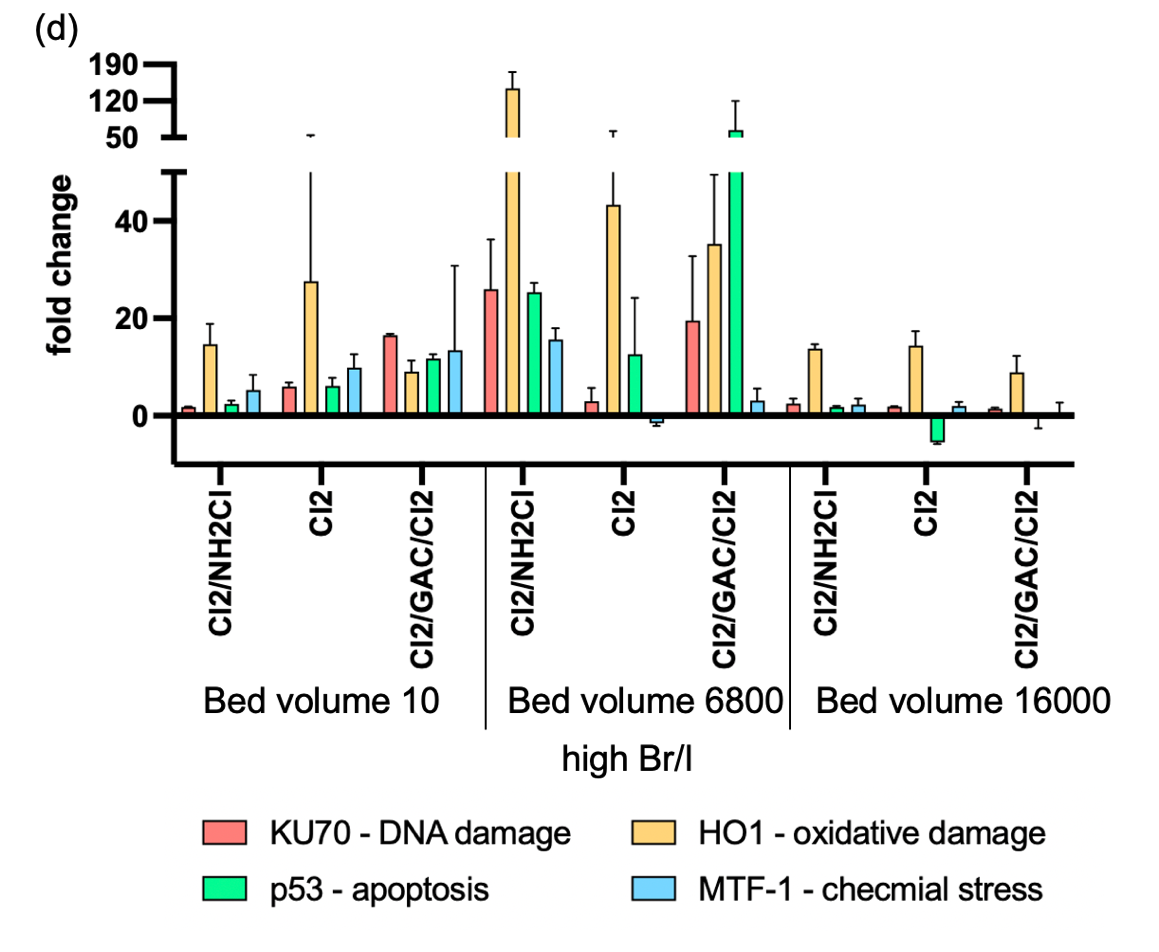

Figure S2: Gene expression fold change in HepG2 cells after 4 hours exposure under of (a) GAC influents treated with free chlorine only (Cl2) and chlorine/chloramines (Cl2/NH2Cl) from GAC system, (b) water samples with low halogen level that were treated by chlorination (Cl2), GAC with post-chlorination (GAC/Cl2), or chlorination/chloramination (Cl2/NH2Cl) from GAC system, (c) water samples with low halogen level that were treated by chlorination (Cl2), GAC with post-chlorination (GAC/Cl2), or GAC with pre- and post-chlorination (Cl2/GAC/Cl2) from GAC system across four concentrations, and (d) water samples with high halogen level that were treated by chlorination (Cl2), chlorine/chloramines (Cl2/NH2Cl), or GAC with pre- and post-chlorination (Cl2/GAC/Cl2) from GAC system at concentration factor of 100. X-axis: fold change of different genes. Y-axis: treatment process.

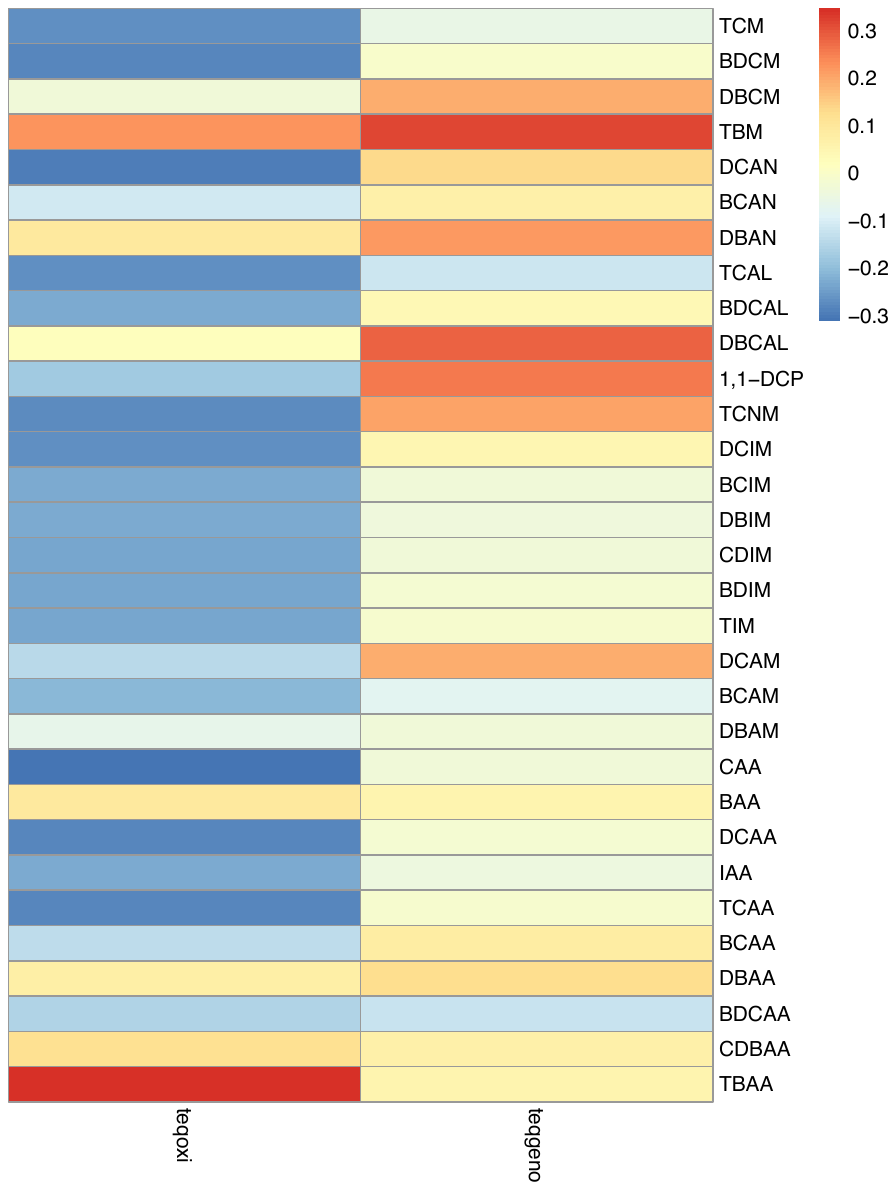

Figure S3: Pearson correlation analysis between DBP concentrations and toxicity endpoint of oxidative stress and DNA stress (TEQ-oxi and TEQ-geno) values in drinking water sample extracts. Correlation coefficients at 95% significance level are scaled with color spectrum on the right, zeros are assigned to the non-significant correlations. Asterisks indicating a significant correlations
